# Regression-guided computational design of auxetic scaffolds for soft tissue applications

**DOI:** 10.1101/2025.08.19.671128

**Authors:** Óscar Lecina-Tejero, Jesús Asín, Jesús Cuartero, María Ángeles Pérez, Carlos Borau

## Abstract

The mechanical performance of tissue-engineered scaffolds plays a critical role in their effectiveness for regenerative medicine applications. Auxetic metamaterials, characterized by a negative Poisson’s ratio, offer enhanced conformability and tunable mechanical behavior, making them promising candidates for scaffold design. This study presents a computational framework combining finite element method (FEM) simulations with regression-based predictive modeling to optimize auxetic scaffold architectures. A design of experiments (DOE) strategy enabled the training of FEM-accurate regression models capable of predicting mechanical responses from scaffold microstructural parameters. Sensitivity analysis guided the development of robust optimization strategies for identifying optimal geometries. Validation of the predictive framework was performed using experimentally derived, skin-representative mechanical properties from published literature, demonstrating strong agreement with FEM results. To facilitate broader use, we developed a software tool integrating this pipeline. It includes a manual mode for direct input of design and geometry parameters, and a predictive mode that returns optimized scaffold designs based on target properties. This integrated methodology supports robust, customizable scaffold design, advancing patient-specific approaches in soft tissue engineering.

## 1 Introduction

Tissue engineering and advanced manufacturing techniques have revolutionized wound healing by developing advanced constructs that mimic the properties of the native tissue extracellular matrix, enabling improved cell proliferation and the regeneration of more complex tissue structures[1]. While traditional tissue engineering scaffolds face challenges in mechanical adaptability, auxetic metamaterials emerge as a promising solution due to their enhanced conformability to dynamic mechanical environments and tunable, design-induced mechanical properties[2, 3].

Auxetic metamaterials are characterized by a negative Poisson’s ratio (NPR), meaning they expand laterally when stretched. This unique property allows them to accommodate significant biaxial deformation while maintaining stable internal stress and adapting their morphology effectively to irregular surfaces[4]. These metamaterials have shown effective mechanical support for tissue engineering and replicated the mechanical behavior of natural soft tissues, which, in addition to their biocompatibility and the improvement of key biological processes such as cell proliferation, migration, and viability, have promoted their use in multiple soft tissue engineering applications[5–8]. Mirani et al.[9] proposed a combined approach of computational modeling and 3D printing to fabricate tissue engineering scaffolds with auxetic designs mimicking the mechanical behavior of porcine pericardium and cardiac valves, while Chansoria et al.[10] investigated the production of hydrogel patches with auxetic geometries adapted to the behavior of dynamic organs, such as lungs, heart, or skin. In the context of soft tissue engineering, understanding the mechanical behavior of the native tissue is essential for designing effective biomimetic scaffolds. For example, human skin exhibits a non-linear and anisotropic behavior influenced by multiple factors including gender, age, body region, thickness, or collagen fiber orientation[11–16]. Figure 1, adapted from the research of Annaidh et al.[13], illustrates these variations. In their study, samples of human back skin were extracted from different regions of the back and aligned distinctly relative to their local Langer lines. The mechanical characterization of these samples revealed varying mechanical behaviors depending on both location and orientation.

**Figure 1:**
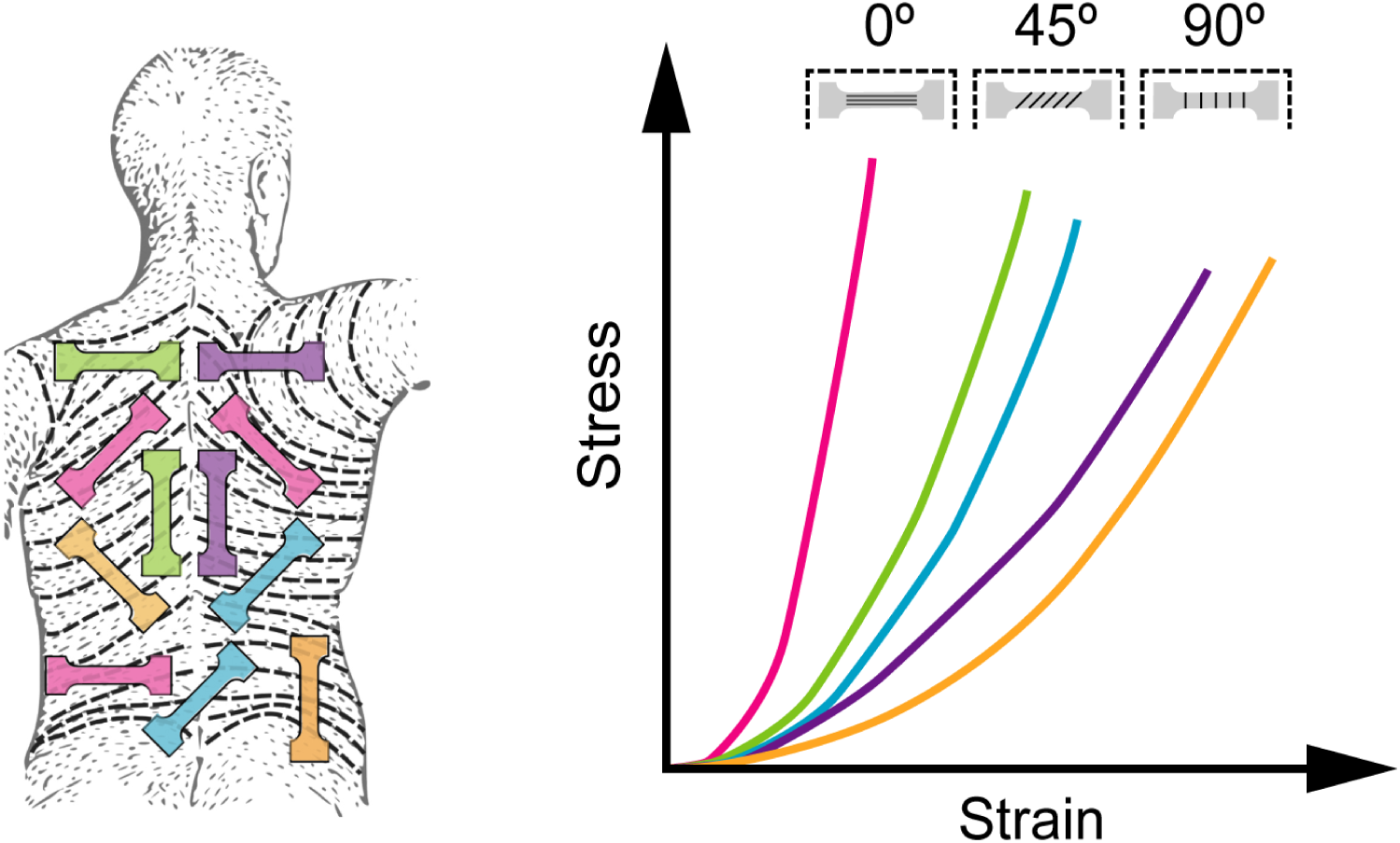
Anatomical location of the skin samples extracted from human back and representative stress-strain curves resulting from mechanical characterization grouped by skin sample orientation with respect to the Langer lines. Figure adapted from [13].

Importantly, the mechanical performance of auxetic metamaterials depends on their design. By controlling the parameters of their microstructural geometry, it is possible to tailor the mechanical properties of the scaffolds to specific functional requirements[2]. Computational analysis using the finite element method (FEM) is a widely adopted tool for structural analysis and design optimization[9], as it enables testing multiple configurations involving geometry, material properties, assemblies, and boundary conditions[17–19]. This allows evaluation of structural performance prior to fabrication, saving time, material, and economic resources. How-ever, FEM analysis of intricate and complex models can incur substantial computational costs, particularly when exploring extensive design spaces with numerous configurations[10, 19]. This increased computational demand may slow the design process and complicate achieving precise mechanical targets. The combination of computational modeling and statistical regression is a validated approach to address these challenges[9], where experimentally validated data can be utilized to train regression models, explore multiple output predictions, and efficiently optimize the design to achieve accurate, prescribed mechanics.

Our work proposed an integrated predictive framework combining FEM simulations, statistical regression modeling, and computational optimization. By employing a design of experiments (DOE) strategy, a reduced number of scaffold configurations can be utilized to generate a representative dataset of FEM computed results. These type of simulations have been validated in a previous work[20]. Regression models trained on this dataset can predict mechanical properties directly from the microstructural design parameters, decreasing the computational effort needed to explore extensive design spaces. Using experimentally obtained human skin mechanical properties (specifically, measurements from back skin samples characterized by Annaidh et al.[13]) as representative targets, this methodology enables the efficient identification of optimal auxetic microfibrous scaffold geometries tailored to patient-specific mechanical conditions. This approach accelerates scaffold development while maintaining accuracy, making it applicable to various soft tissue engineering applications. To support practical implementation, we provide a computational tool[21] that enables users to select optimal design parameters to achieve specific mechanical behaviors.

## 2 Materials and Methods

This study follows an integrated computational workflow designed to predict and optimize the mechanical behavior of auxetic fiber-based scaffolds for soft tissue engineering. The process begins by defining four auxetic geometries, each parameterized to allow systematic variation in scaffold design. A DOE approach was employed to generate a representative set of scaffold configurations by varying key geometric parameters. FEM simulations were then performed on these configurations to assess their mechanical behavior under biaxial loading conditions. The resulting data were used to train statistical models capable of predicting scaffold effective stiffness and strain as a function of design parameters. Finally, optimization techniques were evaluated and applied to identify scaffold geometries that best match specific mechanical targets. This framework enables the rapid and reliable selection of scaffold designs tailored to desired mechanical performance.

### 2.1 Selected auxetic geometries

We have validated our simulations[20] with actual scaffolds fabricated via melt electro-writing (MEW), a technique that provides precise control over microfiber deposition during the printing process. This precision allows for the implementation of specific printing paths to create intricate fibrous microstructures. Consequently, all scaffold geometries presented in this work are specifically designed for fabrication by MEW[22–24]. We selected four re-entrant auxetic designs to define the microstructure of the scaffolds based on their suitability for MEW fabrication, designated as *H-cell* (*HCELL*), *S-regular* (*SREG*), *S-inverted* (*SINV*), and *S-triangular* (*STRI*). These designs were characterized by basic unit cells and parameterized as illustrated in Figure 2 to investigate the influence of geometric variations in auxetic designs on the mechanical response of the scaffolds. The parameters “*a*” and “*b*” were defined to capture geometric variations in fiber layout, while the parameter “*d*” controlled the fiber diameter. The parameters “*xr*”, “*yr*”, and “*zr*” defined the scaffold’s physical dimensions (i.e., the number of unit cell repetitions in the *x* and *y* directions, and the number of stacked layers in the *z* direction).

**Figure 2:**
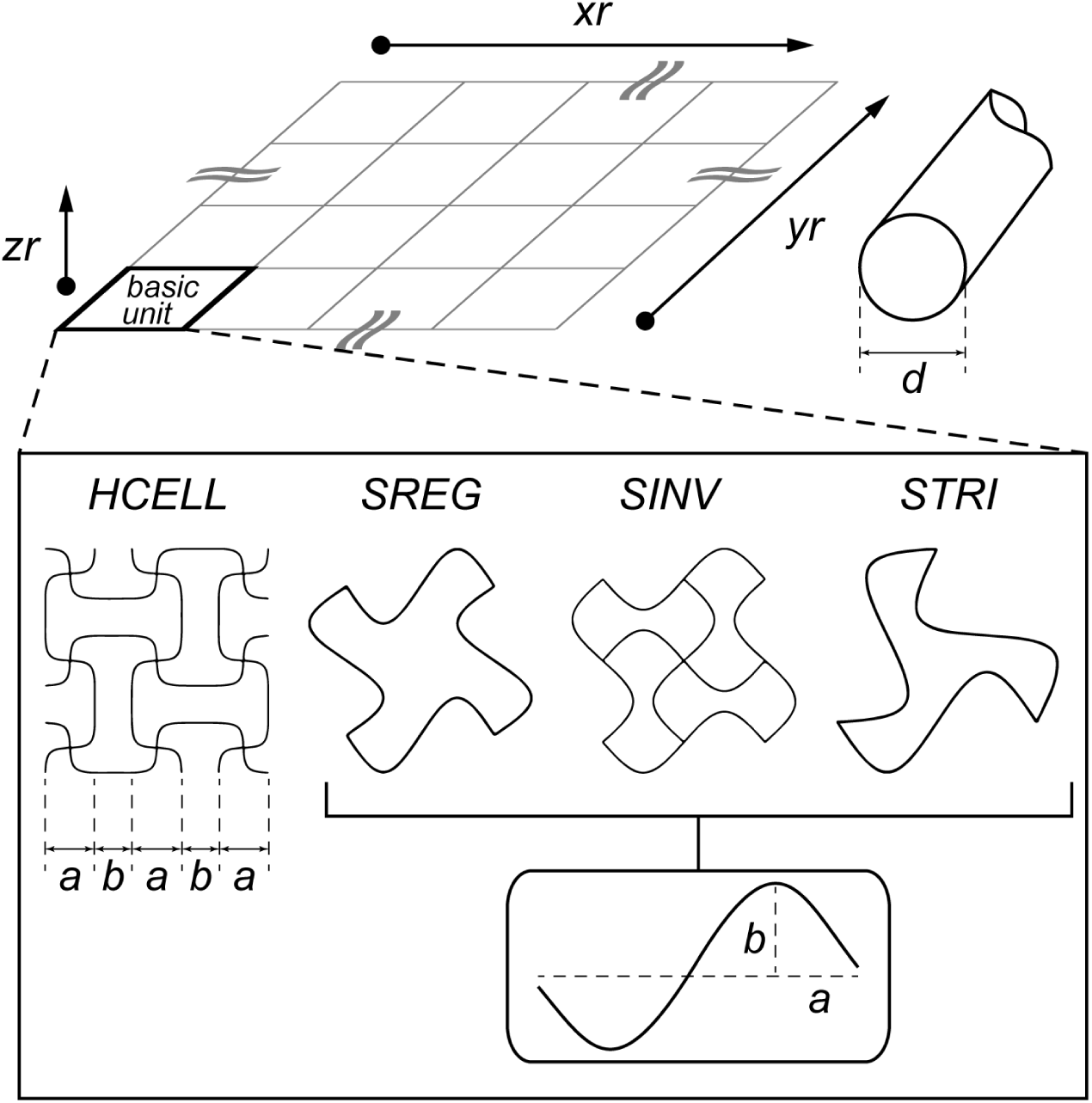
Scheme of auxetic geometry designs and representation of the geometry configurations and parameters. “a” and “b” are directly related to the specific geometries of each auxetic design. Parameter “d” represents the fiber diameter while “xr”, “yr”, and “zr” represent the number of repetitions of the basic unit in these directions, constituting the scaffold volume.

### 2.2 Design of experiments

The design space was explored using a DOE strategy by defining three levels (lower, medium, and higher) for the selected design parameters, which were “*a*”, “*b*”, “*d*”, and “*yr*”. Among them, parameter “*a*” played a central role, as it determined the overall scale of the auxetic geometry and, consequently, the size of the scaffold. To ensure a reduced scaffold scale suitable for tissue engineering applications, its values were constrained between 100 and 300 microns. Smaller “*a*” values may lead to fabrication challenges and compromised structural fidelity, while larger values could result in oversized features unsuitable for scaffold considerations.

It is important to note that parameter “*b*” was defined relative to parameter “*a*” to address design scalability and to comply with the geometric constraints of each auxetic design. Specifically, for the *SREG* and *STRI* designs, parameter “*b*” was established as 0.5, 1, and 1.5 times the value of “*a*”, since higher ratios would lead to fiber overlaps and distortions that compromise the auxetic effect, and smaller values would result in almost straight fibers, not demonstrating any auxetic performance. Therefore, 1.5 was considered the upper limit for the “*b*” to “*a*” ratio (“*rba*”, Eq. 1) across the designs. For the *SINV* design, the same geometric constraints applied, but in this case it restricted the parameter “*b*” to not exceed “*a*”. As a result, the “*rba*” ratios investigated for this design were limited to 0.25, 0.5, and 0.75. In contrast, the *HCELL* design imposed no such restrictions on this ratio, allowing the use of the same range applied in the *SREG* and *STRI* designs.

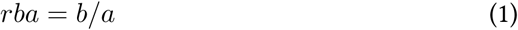

Parameters “*d*” and “*yr*” did not present design-specific limitations and were varied uniformly across all four designs. Specifically, “*d*” was evaluated between 10 and 20 microns, which are common fiber diameters obtained with MEW[20], and parameter “*yr*” ranged from 3 to 9 unit cell repetitions. Parameters “*xr*” and “*zr*” were fixed at values of 6 and 10, respectively, so their combination with the varying values of “*yr*” allowed exploration of the influence of scaffold shape aspect ratio on the mechanical behavior, while also reducing the number of required configurations.

This parameter combination (“*a*”, “*rba*”, “*d*”, “*yr*”) yielded a DOE dataset comprising 324 unique scaffold configurations, with 81 configurations per auxetic design. Table 1 summarizes the parameter values explored for the four auxetic designs.

**Table 1:** Summary of the auxetic designs and the values of their geometry parameters considered in the design of experiments. To maintain design consistency, parameter “*rba*” had to take different values for *SINV* design, while remaining unaltered for the other auxetic designs. The values were attributed to the parameters aiming to explore the design space, comprising both reduced scales feasible by MEW-fabrication and wider scales but without excessively increasing the features, which could lead to lose scaffold-like scales in the structures. The combination of all the possible configurations of the four designs followed this formula: *V alues^P^ ^arameters^* ∗ *Designs* = 3^4^ ∗ 4 = 324, constituting a database of 324 distinct scaffold geometry configurations, with each design represented by 81 unique configurations.

| Geometry parameter | Auxetic design | Lower value | Medium value | Higher value |
| --- | --- | --- | --- | --- |
| $a$ [$\mu m$] | *ALL* | 100 | 200 | 300 |
| $rba$ [—] | *HCELL* | 0.5 | 1 | 1.5 |
|  | *SREG* |  |  |  |
|  | *STRI* |  |  |  |
|  | *SINV* | 0.25 | 0.5 | 0.75 |
| $d$ [$\mu m$] | *ALL* | 10 | 15 | 20 |
| $yr$ [—] | *ALL* | 3 | 6 | 9 |

### 2.3 Numerical simulations

The mechanical performance of the scaffolds was evaluated by conducting FEM-based simulations with ABAQUS software (3DS Dassault Systems). The FEM model of the scaffolds, developed and experimentally validated on a previous work[20], replicates their fibrous structure with 3D beam linear elements (B31) with mesh controls over the length-to-diameter ratio of the elements and node coupling to interconnect the adjacent layers. FEM simulations reproduced the mechanical conditions of the scaffolds during biaxial tensile tests based on control over applied displacement and utilizing a quarter-symmetry model of the scaffolds to reduce computational costs. The Python-based ABAQUS scripts utilized to generate the FEM models are publicly available on GitHub[21, 25].

Despite being one of the main characteristics of auxetic metamaterials, Poisson’s ratio was not extracted from the FEM simulations due to the inherent mechanical configuration of the biaxial tensile tests, which applied symmetric loading in both principal directions. Instead, post-processing of the FEM simulation data focused on identifying the point at which scaffold fibers become fully stretched, marking the limit beyond which auxetic behavior is no longer exhibited. At that point, we measured the strain (*ε*_aux_) and calculated the elastic modulus (*E*_aux_). The calculation of these key variables starts by extracting nodal displacements from the stretched edges and reaction forces from the constrained edges, which allowed to obtain the effective stress (*σ* [kPa]) and strain (*ε* [*µm* / *µm*]) of the scaffolds following the procedure explained in Supp. Info (Figure S7). To consistently define the transition from the initial compliant behavior to the stiffer mechanical response, considering the variability of curve shapes, *ε*_aux_ was defined as the strain at which the instantaneous effective elastic modulus (i.e., the first derivative of the *σ*-*ε* curve, *E* [kPa]) reaches approximately half of its maximum value during the ascending phase, as represented in Figure 3-A and Eq. 2. This relative criterion enabled robust identification of the transition point across the scaffold designs in both *x*– and *y*-directions, and the most restricting direction in each case (i.e., the one with the higher *E*_aux_) was selected to characterize the scaffold behavior.

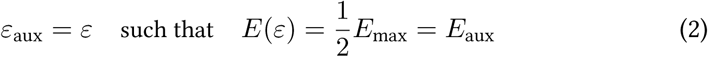

**Figure 3:**
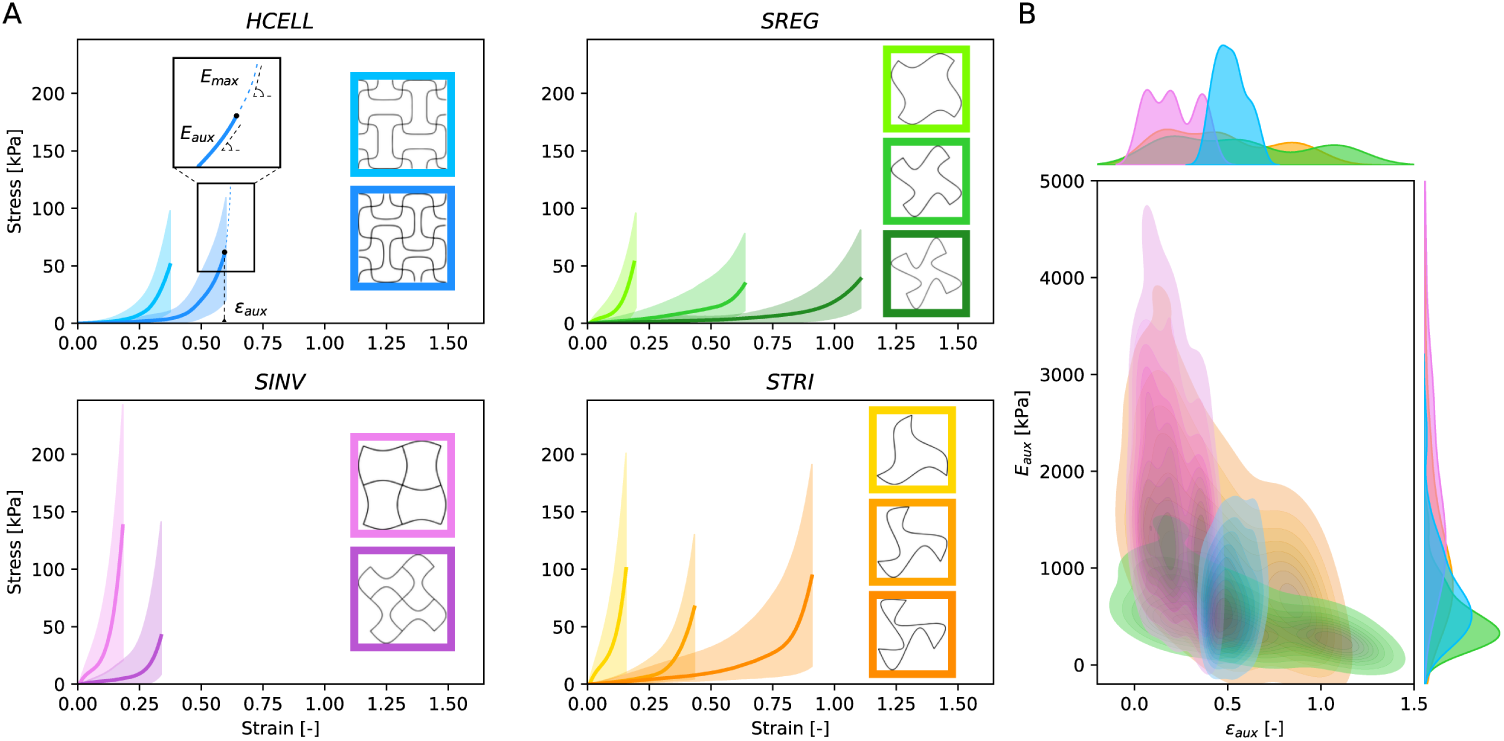
A: FEM-simulated stress-strain curve families for the four auxetic designs and distinct geometric configurations. Curve families were grouped according to observed patterns to enhance the visual interpretation of trends resulting from variations in the geometric configurations of the auxetic designs. B: Distribution of *E_aux_* and *ε_aux_* values, for each auxetic scaffold design (*HCELL*, *SREG*, *SINV*, *STRI*) based on FEM simulation results. The scatter plots represent the *E_aux_* and the *ε_aux_* at the points where FEM simulations became completely extended.

*E*_aux_ and *ε*_aux_ have been defined in alignment with what has been previously described as “Zone 3” in studies of native soft tissues, such as porcine skin[26], where the fibrous components become fully aligned and contribute to a increase in stiffness. This analogy supports the relevance of *E*_aux_ and *ε*_aux_ as representative parameters of the mechanical behavior of the scaffolds in its load-bearing regime. These properties, initially extracted as output responses from FEM simulations, will later serve as key input variables in the operative phase of the study, guiding predictive modeling and scaffold design optimization.

### 2.4 Statistical modeling and optimization procedure

This stage of the methodology addresses the challenge of characterizing the relationship between the scaffold design parameters and their mechanical responses, specifically predicting *E*_aux_ and *ε*_aux_ from geometric variables such as fiber geometric disposition (“*a*” and “*rba*”), fiber diameter (“*d*”), or scaffold aspect ratio (“*yr*”). Despite the extensive DOE performed, the resulting dataset only samples a limited portion of the full parameter space, making it difficult to directly define an explicit function such as Eq. 3:

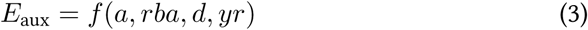

To overcome this limitation, statistical modeling was employed to extract signal from the data and to capture the inherent variability observed in the mechanical response curves. Regression models served two main purposes at this stage. First, they identify which design parameters influence the mechanical behavior of the scaffold, thus enabling dimensionality reduction by eliminating non-influential variables. Second, they provide prediction intervals for the responses, which define a set of possible scaffold configurations compatible with a desired target behavior.

Following the statistical modeling phase, an optimization step becomes necessary to address the inverse design problem: determining which values of the design parameters yield a target mechanical response, represented in Eq. 4:

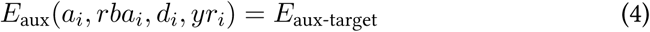

The inverse problem is analytically intractable due to the non-linear and potentially multi-modal nature of the relationships between design parameters and mechanical responses. Therefore, optimization algorithms are required to explore the solution space defined by the statistical model.

The statistical analysis began by fitting regression models relating the FEM simulation outputs (*E*_aux_ and *ε*_aux_) to the geometric parameters of the auxetic designs. For each design, separate regression equations were constructed for each response variable, resulting in a total of eight models. The initial formulation used in all cases included main effects, two-way interaction terms, and quadratic terms, allowing the models to capture a wide range of potential relationships and to represent linear and nonlinear trends, as well as interaction effects. This is represented in Eq. 5:

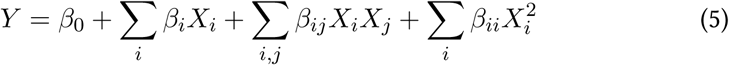

where *Y* represents either *E*_aux_ or *ε*_aux_, depending on the model, and *X_i_*corresponds to the scaffold design parameters (“*a*”, “*rba*”, “*d*”, and “*yr*”). Each auxetic design is modeled independently, allowing the regression equation to reflect design-specific mechanical responses and sensitivities.

Stepwise regression was then conducted in both forward and backward directions to iteratively include or exclude predictor terms based on statistical significance (p-value < 0.05), with the Bayesian Information Criterion (BIC) used to guide model selection. This procedure was independently applied to each response variable (*E*_aux_, *ε*_aux_) for all four auxetic designs, yielding eight distinct regression models.

Once trained, these models functioned as surrogate predictors of the scaffold’s mechanical response based on its geometric parameters. However, prediction alone does not resolve the inverse design problem: determining the input parameters that will yield a specific target mechanical behavior. To address this, an optimization framework was implemented to explore the defined design space for configurations that achieve target values (e.g., *E*_aux-target_) while ensuring robustness against geometric variability.

Several optimization strategies were tested within the DOE-defined space, including random grid search, brute-force exploration, and genetic algorithms. These approaches are well-suited for problems where the parameter space is constrained and sampling is computationally expensive [27]. All optimization strategies aimed to minimize a custom loss function (Eq. 6) that balances two critical aspects of inverse design: *predictive accuracy* and *robustness*. The first term in the equation represents the bias, that is, the mean squared error (MSE) between the average prediction and the target value, ensuring the design meets the desired mechanical response. The second term penalizes high variance among predictions, promoting *reliability* by favoring designs that are less sensitive to small perturbations in geometric parameters. This dual-objective formulation encourages scaffold configurations that are both precise and robust against variability in fabrication or parameter estimation.

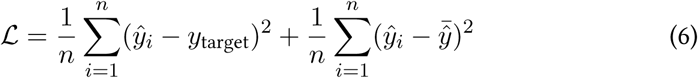

Here, L denotes the loss function to be minimized, *y*^*_i_* are the surrogate model predictions for each sampled geometry, *y*_target_ is the desired target value, and 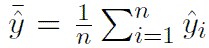 is the mean of the predictions.

Details on the comparative performance of these optimization methods, and the rationale for selecting the final approach, are discussed in the Results section and Supp. Info (Table S2).

R software functions were designed to apply the random grid search method [28, 29], which began with a seed-based sampling of geometric configurations randomly drawn from the DOE space. This seed set was expanded to a pool of up to 5,000 unique configurations, each evaluated using the objective function (Eq. **??**). Sensitivity analysis indicated that the choice of random seed and sample size had minimal impact on the overall outcomes. In contrast, brute-force analysis exhaustively evaluated approximately 770,000 configurations, generated by combining 100 evenly spaces values of “*a*” and “*rba*” with all feasible integer values of “*d*” and “*yr*”. Lastly, a genetic algorithm was implemented using the GA [30] package in R, applying evolutionary operations such as selection, crossover, and mutation to iteratively converge toward optimal solutions within the parameter space.

## 3 Results

### 3.1 FEM simulations demonstrate the design-dependent, non-linear mechanical behavior of auxetic scaffolds

The 324 FEM models of auxetic microfibrous scaffolds were simulated replicating the conditions of biaxial tensile testing described in our previous work[20]. Figure 3-A presents the simulated stress-strain relationships of the four auxetic geometry designs, with the curves grouped according to distinguishable patterns to enhance clarity.

The FEM simulation results reveal that auxetic scaffolds exhibit a non-linear mechanical response, commonly referred to as “J-shaped” behavior, characteristic of soft tissues. This begins with an initial low stiffness (toe region), which progressively increases with stretching (heel region) until reaching a linear region, corresponding to the full expansion and alignment of the scaffold fibers along the loading direction. The *HCELL* and *SINV* designs display two groups of curves (Figure 3-A, left panels), both exhibiting low strain values, indicating a limited deformation capacity. However, the higher stress values observed in the *SINV* curves clearly demonstrate a stiffer mechanical response. In contrast, the *SREG* and *STRI* designs show three groups of curves reaching higher strain values (Figure 3-A, right panels), suggesting greater deformation capacity. Notably, the low stress values associated with the *SREG* design reflect the softest mechanical behavior among the four auxetic geometries. These findings underscore the mechanical versatility of the auxetic scaffolds, where variations in design and geometry can yield a broad spectrum of flexible or resilient mechanical responses.

Both inter-design and intra-design variability of *E*_aux_ and *ε*_aux_ were analyzed to better understand the mechanical properties that auxetic microfibrous scaffolds can exhibit depending on their design and geometric configuration. Figure 3-B shows the kernel density estimations (KDE) of *E*_aux_ and *ε*_aux_ across the four auxetic scaffold designs, highlighting both the overall trends and the variability associated with each design.

The *HCELL* design exhibits a narrow, well-defined distribution in both variables, with *ε*_aux_ values ranging between 0.4 and 0.6, and *E*_aux_ remaining below 1500 kPa. This indicates a consistent mechanical response, suggesting robust and reproducible behavior. In contrast, the *STRI* design displays a broad distribution, reflecting high variability in mechanical performance: *E*_aux_ reaches up to 4000 kPa, while *ε*_aux_ spans from near zero to values exceeding 1. The *SREG* design shows similarly high variability in *ε*_aux_, reaching up to 1.5 (the widest range among all designs) while maintaining a narrower, more concentrated distribution of *E*_aux_ that barely exceeds 1000 kPa. This suggests relatively stable stiffness with greater variability in extensibility. Conversely, the *SINV* design demonstrates the opposite trend, with *E*_aux_ varying widely up to 5000 kPa, and *ε*_aux_ constrained below 0.5, indicating predictable deformation behavior with less predictable stiffness. These observations highlight how the spread and concentration of each design’s mechanical outputs reflect their predictability: narrower distributions imply consistent performance, whereas broader ones indicate greater variability and tunability. The optimal design choice thus depends on the desired trade-off between mechanical stability and adaptability.

Another approach to analyzing the FEM simulation results involved a descriptive exploration of the dataset, focusing on the relationships between geometric parameters (“*a*”, “*rba*”, “*d*”, “*yr*”) and the mechanical outputs (*E*_aux_, *ε*_aux_) for each scaffold design. This analysis, illustrated in Supp. Info (Figures S8–S11), provides deeper insight into how design-specific geometries influence mechanical performance.

In summary, across all four designs, *E*_aux_ consistently exhibits a strong dependency on the predictor variables, though the nature of this dependency varies by design. In the *HCELL* configuration, both *E*_aux_ and *ε*_aux_ display relatively narrow distributions, with moderate sensitivity to all input parameters, suggesting a balanced influence and a robust, predictable mechanical behavior. Notably, increases in “*a*” and “*d*” are associated with reduced/increased stiffness respectively, while “*rba*” exerts a clear inverse influence on strain. The *SREG* design shows a pronounced sensitivity of *ε*_aux_ to the “*rba*” ratio, with strain increasing significantly at higher values, while *E*_aux_ is most affected by “*a*”, “*rba*” and “*d*”, indicating that structural extensibility is more tunable than stiffness. This is consistent with the broader spread observed in strain outputs for this design, reaching values up to 1.5. In the *SINV* design, *ε*_aux_ remains confined below 0.5 regardless of geometric variations, pointing to limited deformability, with “*rba*” being the only parameter with a clear effect on the achieved strain. On the other hand, *E*_aux_ is highly responsive to both “*a*” and “*d*”, and to a less extent to “*rba*”, ranging up to 5000 kPa and making this design suitable for applications requiring tunable stiffness with constrained strain. The *STRI* design exhibits the highest overall variability. *ε*_aux_ again shows a dominant dependence on “*rba*”, with strain values spanning a wide range, while *E*_aux_ responds to a combination of “*a*”, “*d*”, and “*yr*”. This high sensitivity across parameters results in a flexible but less predictable mechanical profile.

An important finding across all designs is the dominant influence of the “*rba*” ratio on the scaffold’s mechanical response, particularly on *ε*_aux_. This parameter, representing the aspect ratio between the primary geometric features of the unit cell, governs the deformation mechanisms underlying auxetic behavior, affecting how the structure expands or contracts laterally under tensile load. By altering the balance between “*a*” and “*b*”, the “*rba*” ratio modulates the kinematic degrees of freedom within the cell, directly impacting both the extent and nature of strain accommodation. Overall, these results highlight how the coupling between design parameters and mechanical responses varies across scaffold architectures. While *HCELL* offers a stable and balanced mechanical response, the other designs, especially *STRI* and *SREG*, enable greater tunability in extensibility, albeit with increased variability.

### 3.2 Statistical modeling and optimization for target mechanical performance

Statistical regression models were built for each of the four auxetic designs to predict the key mechanical outputs (*E*_aux_ and *ε*_aux_) based on geometric input parameters from the DOE database. To evaluate the quality of these models, predictions were compared to FEM simulation results for the corresponding geometric configurations. As shown in Figure 4, the alignment of the scatter points with the diagonal line reflects a strong agreement between predicted and observed values. Among the models, *SINV* demonstrated the highest accuracy, while *STRI* exhibited greater prediction variability, especially at lower *E*_aux_ values. In particular, the *E*_aux_ models exhibited high predictive accuracy across all designs, with Adj. R^2^ values exceeding 0.96. The *ε*_aux_ models showed more variable performance, with higher accuracy for the *HCELL* and *SINV* designs (Adj. R^2^ > 0.99), and lower accuracy for *SREG* and *STRI* (Adj. R^2^ > 0.82).

**Figure 4:**
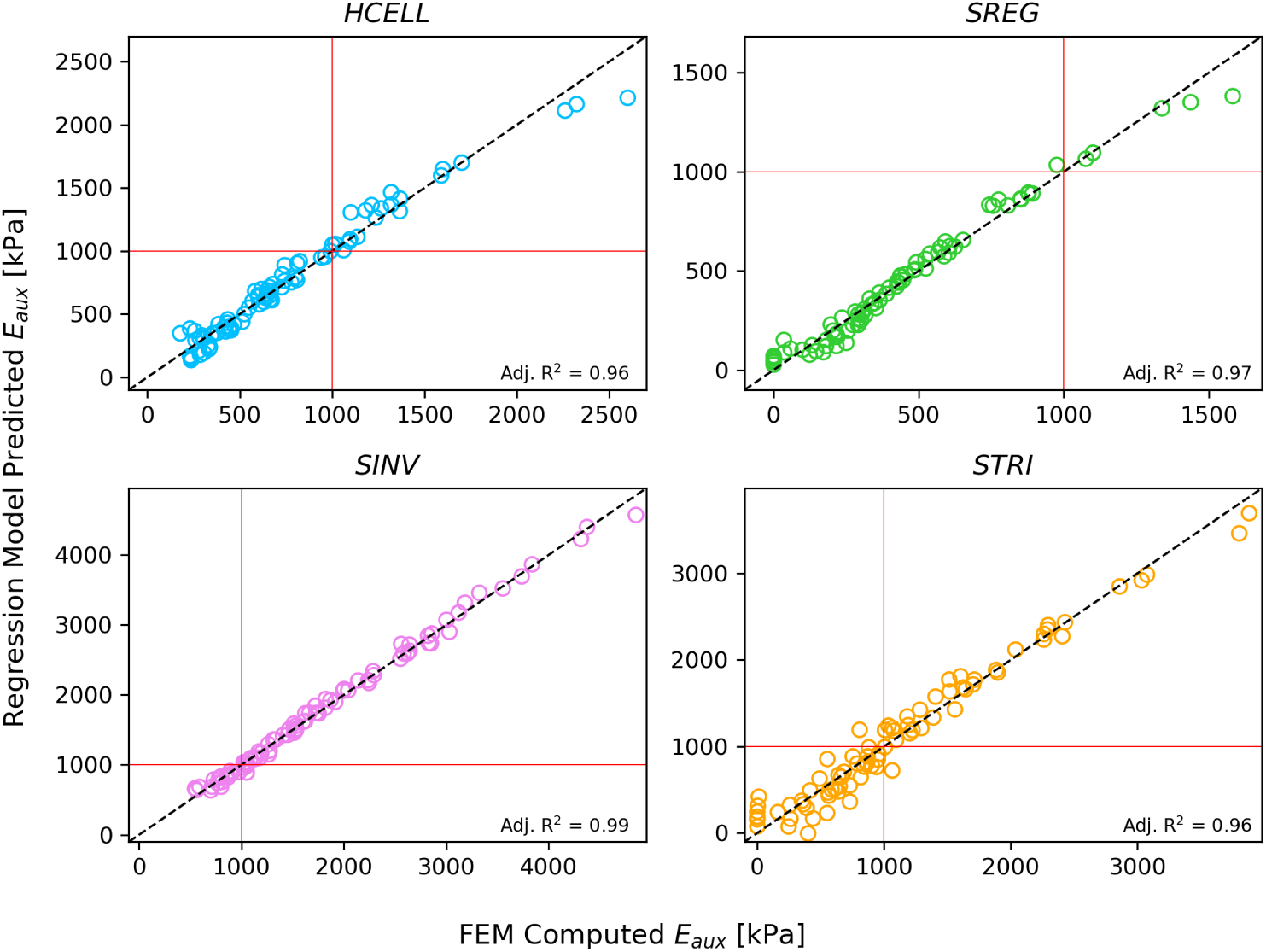
Predicted vs. FEM observed values of *E_aux_* for the four auxetic scaffold designs, demonstrating the accuracy of the regression models, which is represented by the high values of Adj. R^2^ (> 0.96). The red lines indicate the representative target value of 1000 kPa, used as a benchmark for evaluating the design optimization strategy and helping the visualization of *E_aux_* range differences across geometries. For example, the *SREG* design shows predictions predominantly below the target, whereas the *SINV* design exhibits a wide distribution with many configurations exceeding the target, highlighting distinct mechanical potential depending on the design selected.

Figure 4 includes a reference target of 1000 kPa (highlighted by red lines), selected as a representative benchmark across all four scaffold designs. This target lies within the range of output values obtained by the FEM simulations and was used here to neutrally compare the performance of the optimization strategies. By decoupling this initial test from the skin-specific targets addressed later, we ensured a consistent and design-independent basis for evaluating the reliability and efficiency of each method.

To assess the predictive robustness of the regression models near this reference target, a sensitivity analysis was carried out using the *STRI* model, which exhibited the lowest Adj. R^2^ value (0.96) and the widest spread between predicted and observed values (see Figure 4). This made it a suitable candidate for testing the stability of the optimization pipeline. The analysis involved generating predictions and 95% confidence intervals for all configurations in the *STRI* training dataset (Supp. Info – Figure S12). Comparison with the original FEM outputs revealed several parameter combinations with tight prediction agreement, validating the performance of the model even under moderate variability.

Using this 1000 kPa target, we then compared the three previously described optimization strategies: i) random grid search; ii) brute-force search and iii) genetic algorithm. Random grid search emerged as the most efficient method, identifying robust solutions in under 10 seconds with relatively few evaluations. Brute-force search, while exhaustive and accurate, required substantial computational resources. The genetic algorithm delivered comparable performance to random grid search but offered no advantages in terms of solution quality or runtime and was therefore not selected for further use. A detailed comparison of the three strategies in terms of computational cost and predictive robustness is provided in Supp. Info (Table S2).

This preliminary assessment enabled the selection of an efficient optimization method, which was subsequently applied to achieve target values discussed in the next section.

### 3.3 Auxetic scaffolds provide reliable solutions to reproduce the behavior of soft tissues

To demonstrate the applicability of the proposed predictive framework, the regression and optimization approach was used to identify auxetic scaffold configurations capable of replicating the mechanical behavior of soft tissues, particularly human skin. Reference values for skin elasticity were obtained from the literature [13], selecting three representative elastic modulus values corresponding to orientations of 90°, 45°, and 0° relative to the local Langer lines. These values (*E*(90^◦^) = 725 kPa, *E*(45^◦^) = 1210 kPa, and *E*(0^◦^) = 1580 kPa) served as target mechanical properties to be matched by the scaffold designs.

For each representative skin modulus, the regression models were employed to predict scaffold geometric configurations that would not only match the target stiffness values but also exhibit stable mechanical behavior. These optimized configurations were subsequently validated through FEM simulations to assess prediction accuracy. Figure 5 summarizes the results obtained for the three skin stiffness targets across the four auxetic scaffold designs, including the optimized geometric parameters, predicted and simulated values of *E*_aux_, their relative errors to the target, and the corresponding predicted and simulated values of *ε*_aux_.

**Figure 5:**
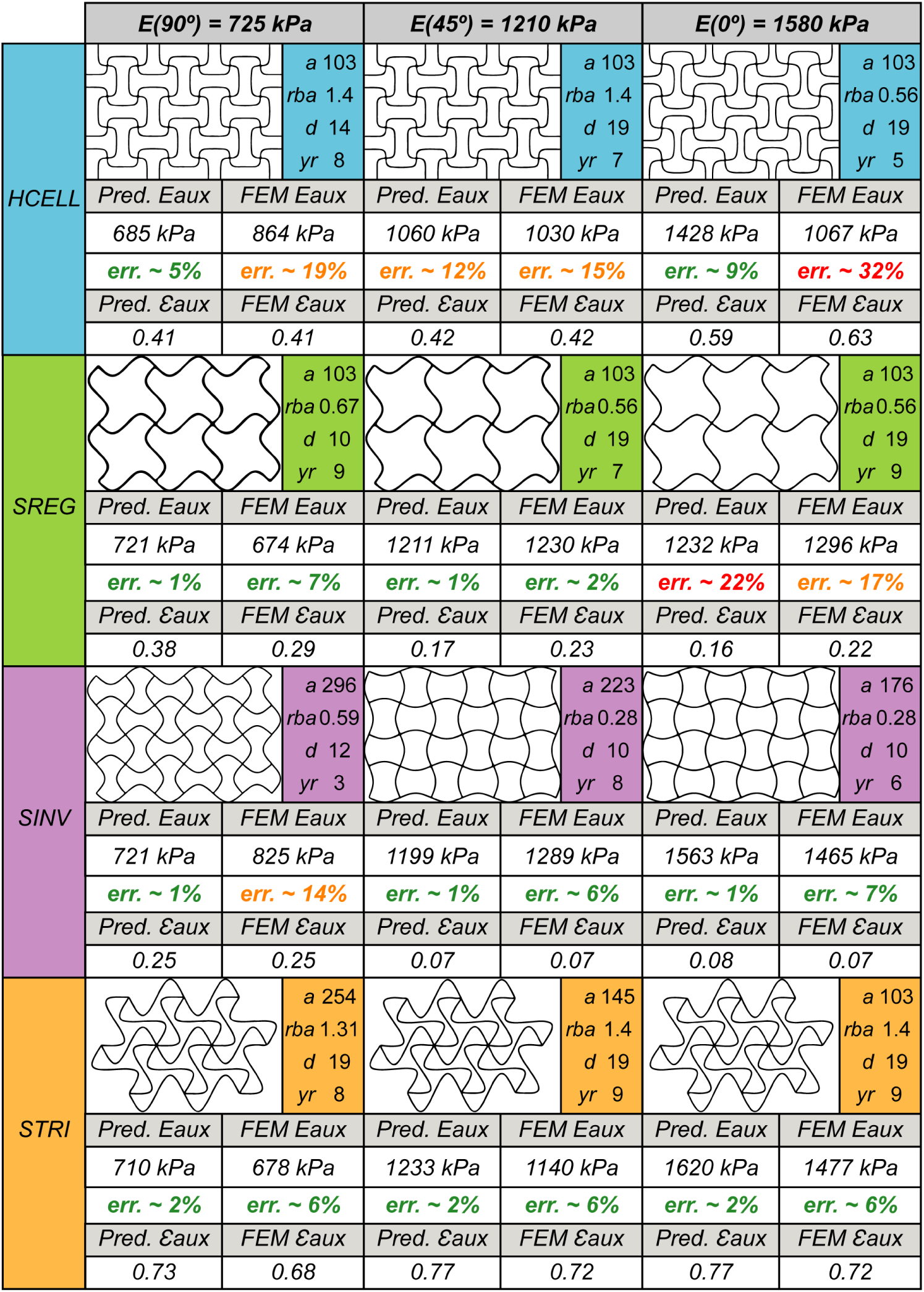
Predictions and FEM results of auxetic scaffolds aiming to match specific skin properties extracted from literature[13]. Predictions were obtained based on each considered target and FEM simulations were conducted to prove the accuracy of the predictions.

The *HCELL* design showed variable performance across the target range. It delivered reasonable accuracy (5%) for the *E*(90^◦^) target, but larger deviations were observed in the FEM validation (19%). For the *E*(45^◦^) target, both predicted and simulated values underestimated the target modulus, though the relative errors remained low (between 12% and 15%). This design struggled to reproduce the higher stiffness value (*E*(0^◦^) = 1580 kPa), with predicted and simulated values falling short (1428 kPa and 1067 kPa, respectively). Nonetheless, predicted and simulated *ε*_aux_ values showed strong agreement across all cases, indicating robust strain estimations.

The *SREG* design achieved excellent accuracy for the lower and intermediate targets (*E*(90^◦^) and *E*(45^◦^)), with errors below 10% in both prediction and simulation. However, it failed to capture the high stiffness target (*E*(0^◦^)), though the consistent correlation between predicted and simulated values across both *E*_aux_ and *ε*_aux_ supports the model’s reliability even when absolute accuracy decreased.

The *SINV* design performed reliably for the intermediate and high targets, providing low-error (between 1% and 7%) predictions and simulations for *E*(45^◦^) and *E*(0^◦^). For the *E*(90^◦^) target, despite an accurate prediction, FEM simulations revealed greater deviation. This design consistently produced lower strain values (around 0.07) than the other scaffolds (between 0.2 and 0.7), though predicted and simulated *ε*_aux_ values remained well-aligned.

The *STRI* design demonstrated the most consistent and accurate performance across all three targets. Prediction errors for *E*_aux_ remained around 2%, with FEM simulation errors under 6%, regardless of the target stiffness. *ε*_aux_ estimations were similarly robust, with close agreement between predicted and simulated values across all configurations.

Overall, these results validate the effectiveness of the predictive framework in identifying scaffold configurations that replicate the anisotropic mechanical behavior of human skin or other tissues. The distinct performance patterns observed across the different scaffold designs indicate that certain architectures are inherently better suited to achieving specific mechanical targets. This ability to guide targeted scaffold selection represents a meaningful advance in the development of biomimetic auxetic scaffolds for skin tissue engineering applications. To support real-world implementation, we developed a computational tool (described in the following section) that automates the selection and generation of printable scaffold geometries based on target mechanical properties

### 3.4 Development of a computational tool for scaffold design optimization and 3D printing

We developed a computational tool[21], including a minimal user interface, that integrates the trained predictive models and optimization algorithms to select optimal parameters and directly generate the corresponding 3D printing or FEM files, streamlining the transition from design to fabrication/simulation and enabling rapid, application-specific scaffold production. This tool allows users to either input specific geometric parameters (Manual Mode) or specify target mechanical properties (Predictive Mode) and automatically obtain the most suitable scaffold geometry.

In the current version, users can input specific values for the geometrical parameters defining the four auxetic designs shown in this work (*HCELL*, *SREG*, *STRI*, and *SINV*) in the “Manual Mode”. Based on these configurations, the platform automatically generates FEM models as ABAQUS input files, G-code files for MEW fabrication, or both. This functionality enables precise control over scaffold design, useful for custom applications or exploratory testing of specific parameter combinations.

In contrast, the “Predictive Mode” encapsulates the core contribution of this study by enabling geometry prediction based on desired mechanical performance. Users are prompted to input a target value for the effective elastic modulus in the toe region, *E*_aux_. The pre-trained regression models are then employed to identify the optimal geometric configurations across each of the four auxetic designs. These configurations are subsequently presented in a comparative format, allowing users to assess and select the most appropriate option. As in the “Manual Mode”, the selected configurations can be immediately exported as FEM input files, G-code files, or both, facilitating direct integration into simulation workflows and 3D printing processes.

A block diagram illustrating the functional steps of the computational tool is shown in Figure 6, outlining the workflow for both modes, by accessing directly to prototype design or starting from mechanical target definition to the generation of design prototypes and exportable files. Supp. Info (Figures S13–S15) includes screenshots of the user interface and example outputs from both design modes, providing a visual reference of the platform in operation.

**Figure 6:**
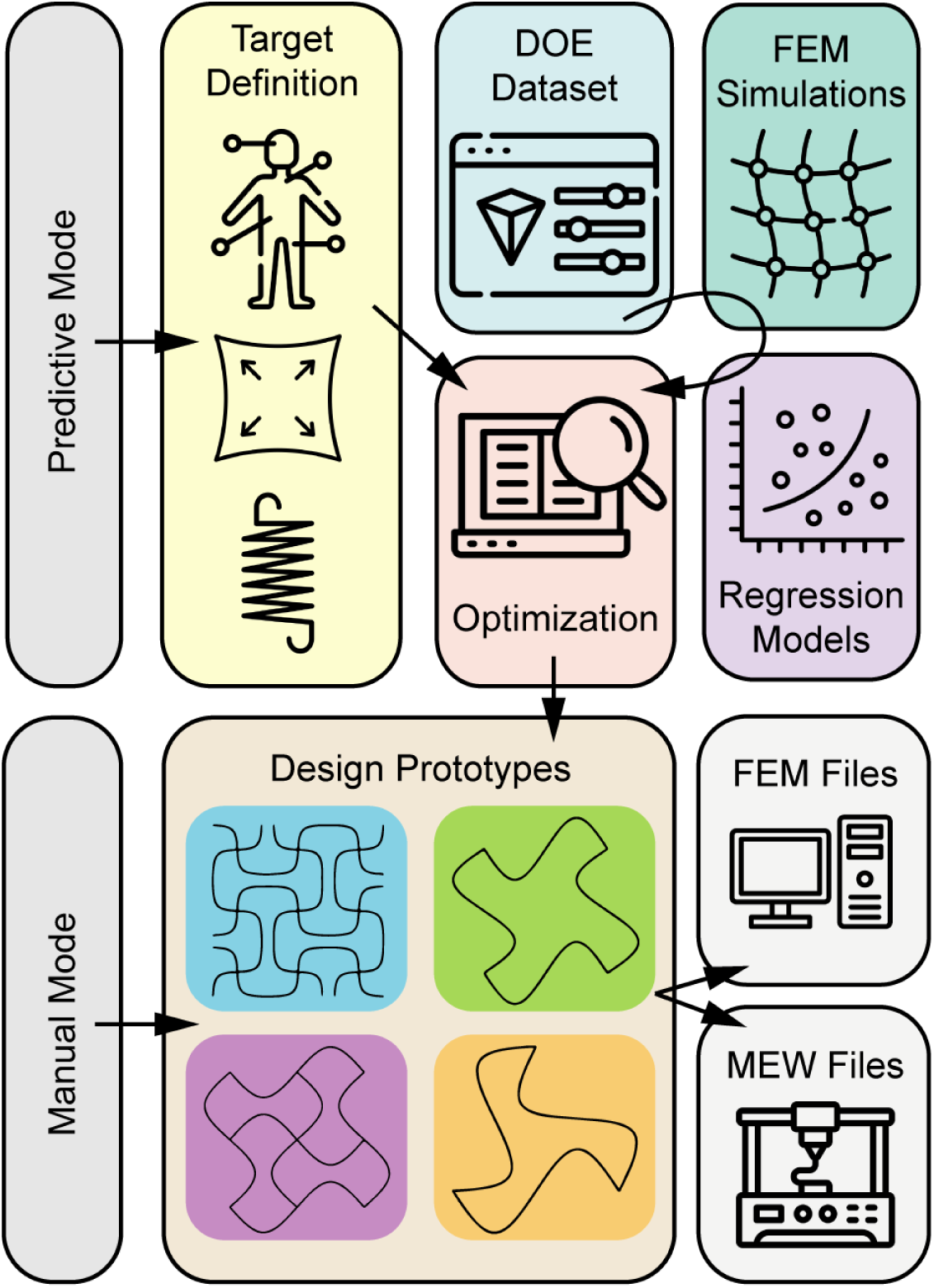
Block diagram of the computational tool workflow. In the “Manual Mode”, the user can access directly to geometric parameter definition and generate the desired configurations, while in the “Predictive Mode” the user is allowed to define a mechanical target. This user-defined target is then fed to the optimization module, which selects the optimal combination of geometric parameters for each of the four auxetic scaffold designs. The optimization process is based on the statistical regression models developed from training with the DOE dataset of FEM simulations, enabling accurate predictions of mechanical responses based on design parameters. This framework allows automated selection of scaffold designs that best match user-defined targets, streamlining the transition from design concept to fabrication.

## 4 Discussion

This study presents a robust computational framework for designing auxetic microfibrous scaffolds with tailored mechanical properties that replicate the anisotropic behavior of human skin and other soft tissues. By integrating finite element simulations with regression-based predictive modeling and optimization techniques, we established an efficient and reliable methodology for identifying scaffold geometries that meet specific mechanical targets. This approach significantly reduces the computational burden typically associated with design optimization, providing a streamlined and accessible solution for the development of mechanically customized scaffolds

The mechanical characterization of the four auxetic designs (*HCELL*, *SREG*, *SINV*, and *STRI*) revealed distinct behavior patterns that align with the fundamental requirements for skin tissue engineering. All designs exhibited the non-linear “J-curve” stress-strain behavior characteristic of soft tissues, with varying degrees of stiffness and deformation capacity (Figure 3). This biomimetic mechanical response is particularly valuable for tissue engineering as it replicates the toe region observed in natural skin, where an initial soft behavior transitions to increased stiffness under loading, which is a feature crucial for proper soft tissue function. The remarkable diversity in mechanical performance across the different auxetic designs offers considerable flexibility in addressing the heterogeneous properties of human skin[13]. *SINV* scaffolds demonstrated higher stiffness with limited deformation capacity, making them suitable for applications requiring greater mechanical resilience such as back, chest, or forearms, where skin has stiffer mechanical properties to serve protective functions [13–15, 31]. In contrast, *SREG* designs exhibited the softest mechanical response with extended deformation capacity, potentially beneficial for more compliant skin regions such as joints or abdomen, which need looser behavior to accommodate frequent movement [31–33]. The *HCELL* and *STRI* designs presented intermediate mechanical behaviors with different degrees of reproducibility and predictability[22].

The statistical distribution of *E_aux_* and *ε*_aux_ values across the design space provides important insights into the reliability and versatility of each geometry. Designs with more compact distributions, such as *HCELL*, offer predictable mechanical responses with reduced sensitivity to geometric variations, which is a valuable characteristic for manufacturing reproducibility. Conversely, designs with broader distributions, particularly *SINV* and *STRI*, provide greater flexibility in achieving diverse mechanical values, although with potentially reduced reproducibility. This exchange between consistency and range of achievable properties must be carefully considered when selecting appropriate designs for specific applications.

The high predictive accuracy of our regression models (Adj. R^2^ > 0.96 for *E*_aux_ and Adj. R^2^ > 0.82 for *ε*_aux_) confirms the effectiveness of the proposed approach in reducing reliance on time-intensive FEM simulations during the scaffold design process. Comparison between model predictions and FEM outputs further validates the framework’s ability to reliably identify optimal scaffold configurations across diverse mechanical targets. Among the optimization techniques tested, random grid sampling emerged as the most efficient, consistently yielding robust solutions with minimal computational overhead. Consequently, this was the method integrated into the optimization core of the computational tool.

When applying our framework to mimic the properties of human skin, we observed varying degrees of success across the different auxetic designs. The *STRI* design demonstrated the most consistent performance across all three skin representative targets (E(90°), E(45°), and E(0°)), with prediction errors around 2% and FEM validation errors around 6%. This consistency indicates the robust adaptability of this particular design across a wide range of stiffness values. The *SINV* design showed excellent performance for intermediate and higher stiffness targets, while *SREG* excelled at lower and intermediate stiffness values. These complementary strengths across different auxetic designs expand the available resources for addressing the diverse mechanical requirements of skin tissue engineering.

The inability of certain designs to accurately reproduce specific target values, particularly the higher stiffness E(0°) target for *HCELL* and *SREG* designs, highlights the inherent limitations of each geometry. This suggests that no single auxetic design can address the full spectrum of skin mechanical properties, emphasizing the importance of a design selection strategy based on specific application requirements. The integration of multiple designs within a single scaffold could potentially provide a more comprehensive mimicry of the complex mechanical behavior of human skin, though this would introduce additional manufacturing challenges.

Our findings align with previous studies that have demonstrated the potential of auxetic materials in tissue engineering applications [2, 3, 5–10]. However, our work extends beyond existing literature by establishing a systematic and accessible methodology for predicting and optimizing scaffold properties, and by incorporating it into a practical and user friendly computational tool.

Nevertheless, several limitations should be acknowledged in the current study. First, our framework focuses primarily on the toe and heel regions of the stress-strain curve, characterized by *E*_aux_ and *ε*_aux_. While this region is particularly relevant for capturing the physiological response of skin under normal loading conditions [26], future studies should extend the analysis to include the full non-linear behavior at higher strain levels, especially in applications where tissues are subject to larger deformations.

Second, the current model considers only static mechanical behavior. However, native skin is subjected to dynamic and time-dependent mechanical loading *in vivo*. Incorporating viscoelasticity, time-dependent effects, and fatigue analysis into the predictive framework would provide a more comprehensive and biologically relevant understanding of scaffold performance under real-world physiological conditions.

Third, our analysis is limited to only four auxetic scaffold designs. While these designs were selected to span a range of geometrical features, the design space for auxetic microarchitectures is vast. Expanding the library of tested geometries and integrating more complex or hierarchical structures could unlock novel mechanical behaviors and improve the framework’s adaptability across different target tissues. Notably, the developed user interface can serve as a flexible platform for future integration of additional geometries, design rules, or optimization algorithms, enhancing its utility as the methodology evolves.

From a manufacturing perspective, while MEW offers precise control over microfiber deposition, practical fabrication constraints may introduce geometric variations that could influence the scaffold’s mechanical performance. Incorporating geometric tolerances into the optimization process would improve the robustness of predicted configurations. In our previous work[20], we explored strategies to minimize these deviations and enhance print fidelity, which could be integrated into future iterations of this computational workflow and reflected in the interface output.

Finally, although this study demonstrates that auxetic scaffolds can be engineered to match target mechanical properties, the successful translation into functional tissue constructs will depend on biological performance. Factors such as cell attachment, proliferation, and extracellular matrix deposition must also be considered. Preliminary studies in our group have shown that these scaffolds support favorable cell behavior under *in vitro* conditions, but more comprehensive biological validation is needed to assess long-term viability and integration in a physiological environment[20].

## 5 Conclusions

This study presents a computational framework that significantly advances the design of auxetic, biomimetic microfibrous scaffolds for skin tissue engineering. By integrating FEM simulations with regression-based predictive modeling and optimization strategies, we established a methodology capable of efficiently identifying scaffold geometries tailored to specific mechanical targets. The distinct performance of each auxetic design across the stiffness spectrum highlights the versatility of the framework in addressing the heterogeneous and anisotropic properties of human skin.

Beyond the methodological advances, we translated this framework into a user-friendly interface that enables both manual design and predictive design based on mechanical targets. This platform streamlines the entire process, from specific mechanical requirements to exporting fabrication-ready files, offering a practical tool for researchers aiming to develop customized auxetic scaffolds.

This approach not only reduces computational demands but also provides a systematic and accessible path for scaffold design based on desired mechanical outcomes. Future work should focus on the experimental validation of the optimized configurations, in-depth investigation of cell-scaffold interactions, and the incorporation of additional factors such as dynamic loading, viscoelastic behavior, and biological cues relevant to skin tissue regeneration.

## 6 Acknowledgements

Authors would like to acknowledge the Spanish Ministry of Economy and Competitiveness through the project PID2023-146072OB-I00.

## 8 Supplementary Material

### 8.1 FEM simulation data post-processing

The post-processing of the FEM simulation data focused on extracting nodal displacements (Δ*u* [*µm*]) from the stretched edges and reaction forces (*RF* [mN]) from the constrained edges, which allowed the calculation of effective stress (*σ* [kPa]) and strain (*ε* [*µm/µm*]) of the scaffolds following Eq. 7 and Eq. 8.

**Figure 7:**
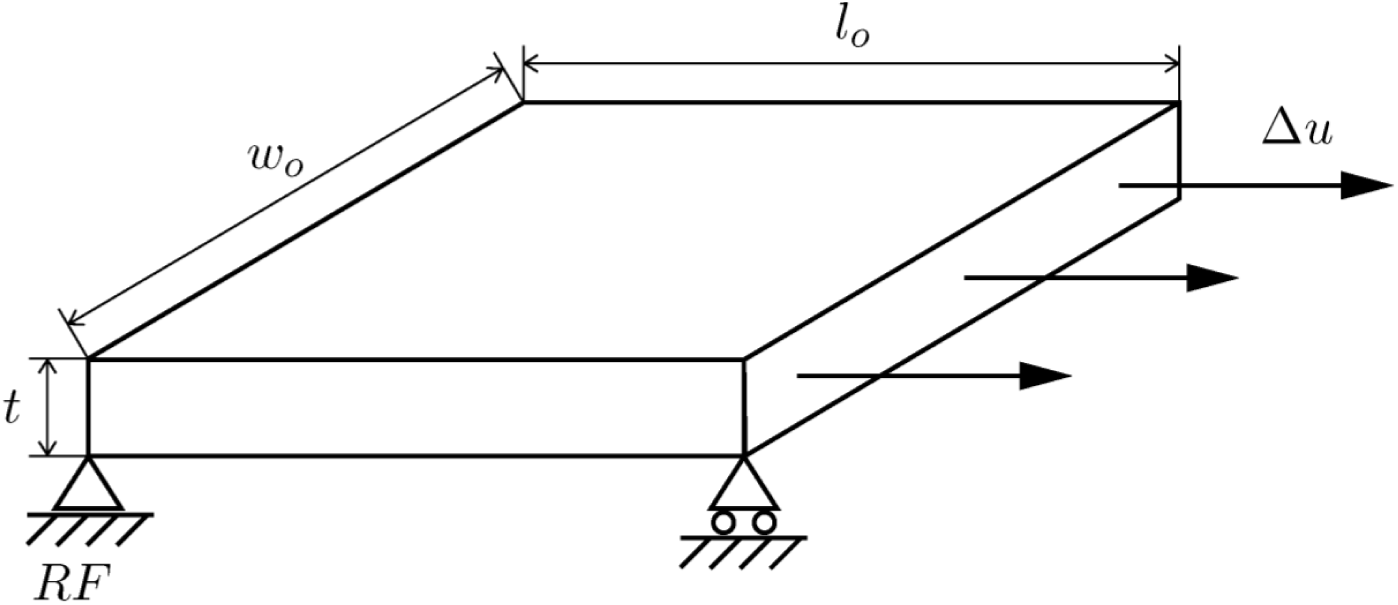
Schematic representation of the dimensions considered for scaffold mechanical properties calculation.

The scaffolds were treated as a homogeneous continuum, with their entire volume considered for length (*l_o_*[*µm*]), width (*w_o_* [*µm*]), thickness (*t* [*µm*]), and the consequent transversal area (*A_t_* [*µm*^2^]) determinations (Eq. 9), as illustrated in Figure 7.

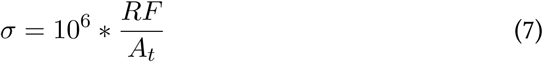

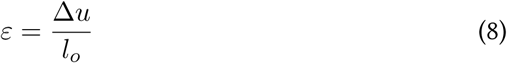

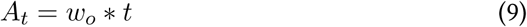

**Figure 8:**
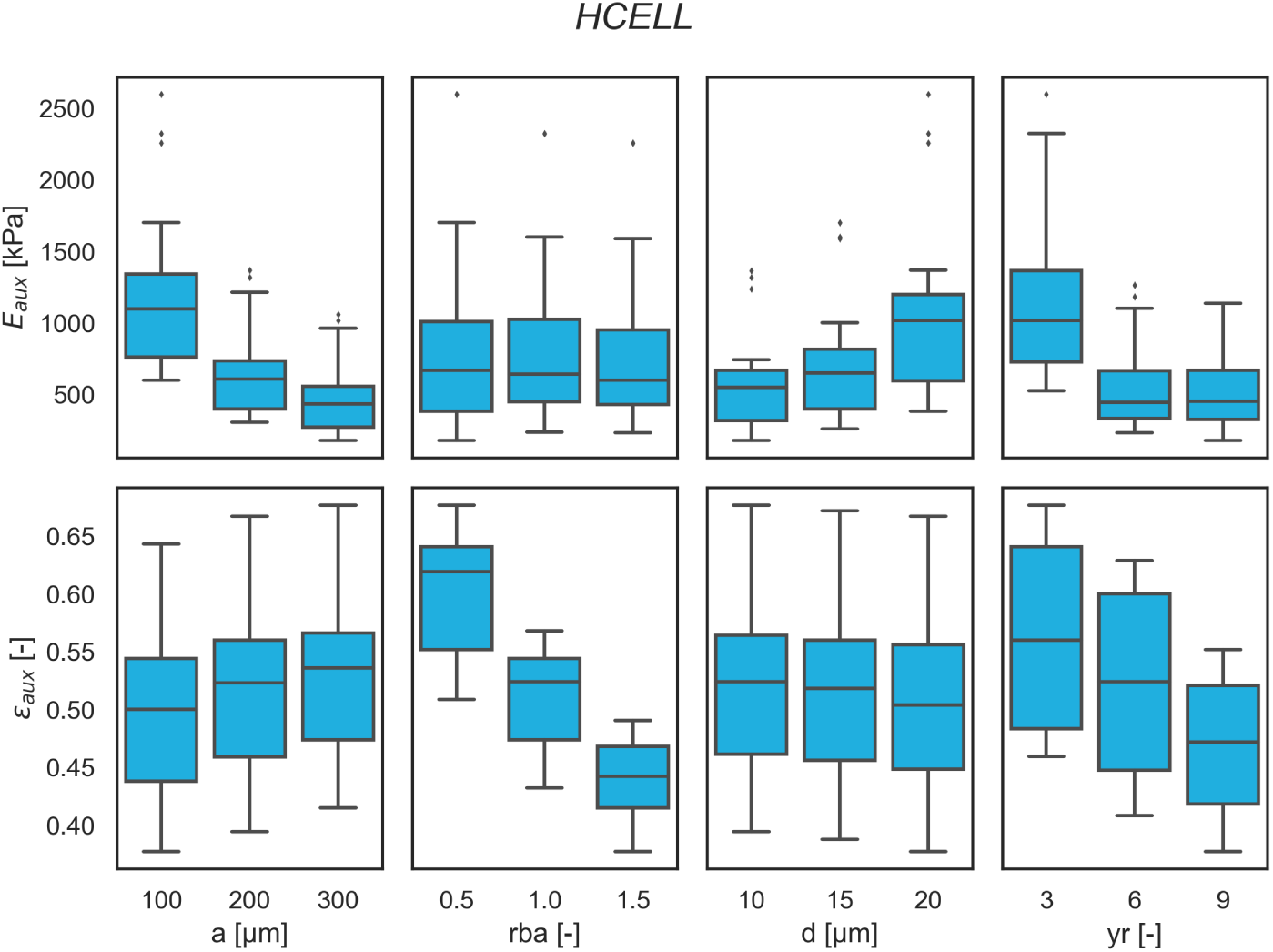
Interactions between geometric parameters and scaffold mechanical response variables observed from *HCELL* design FEM simulations. This design exhibits balanced sensitivity to all parameters, resulting in a stable and predictable mechanical response with moderate tunability in both stiffness and strain.

**Figure 9:**
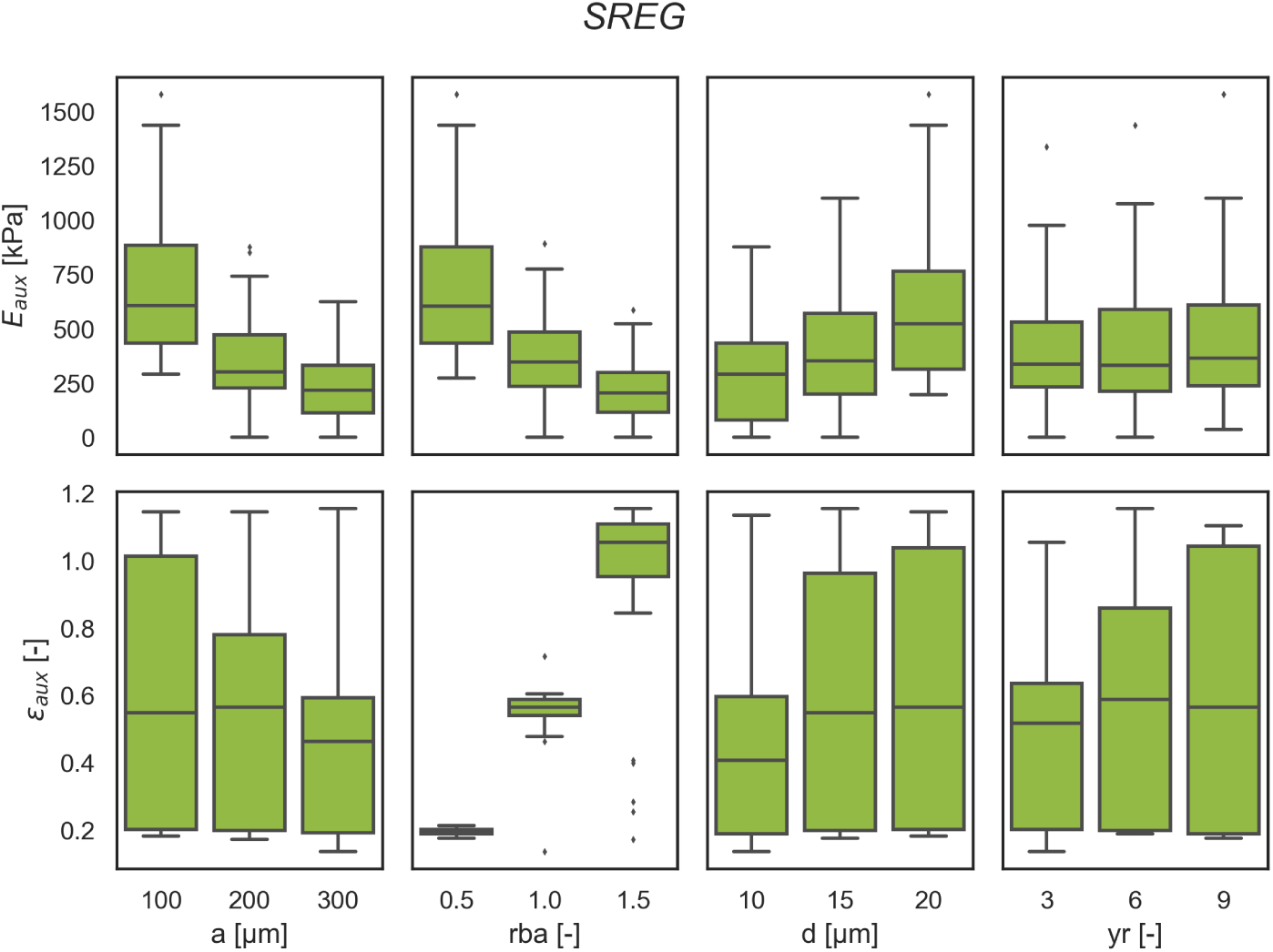
Interactions between geometric parameters and scaffold mechanical response variables observed from *SREG* design FEM simulations. Characterized by high strain variability and strong dependence on the “*rba*” ratio, *SREG* enables significant extensibility at the cost of increased mechanical unpredictability.

**Figure 10:**
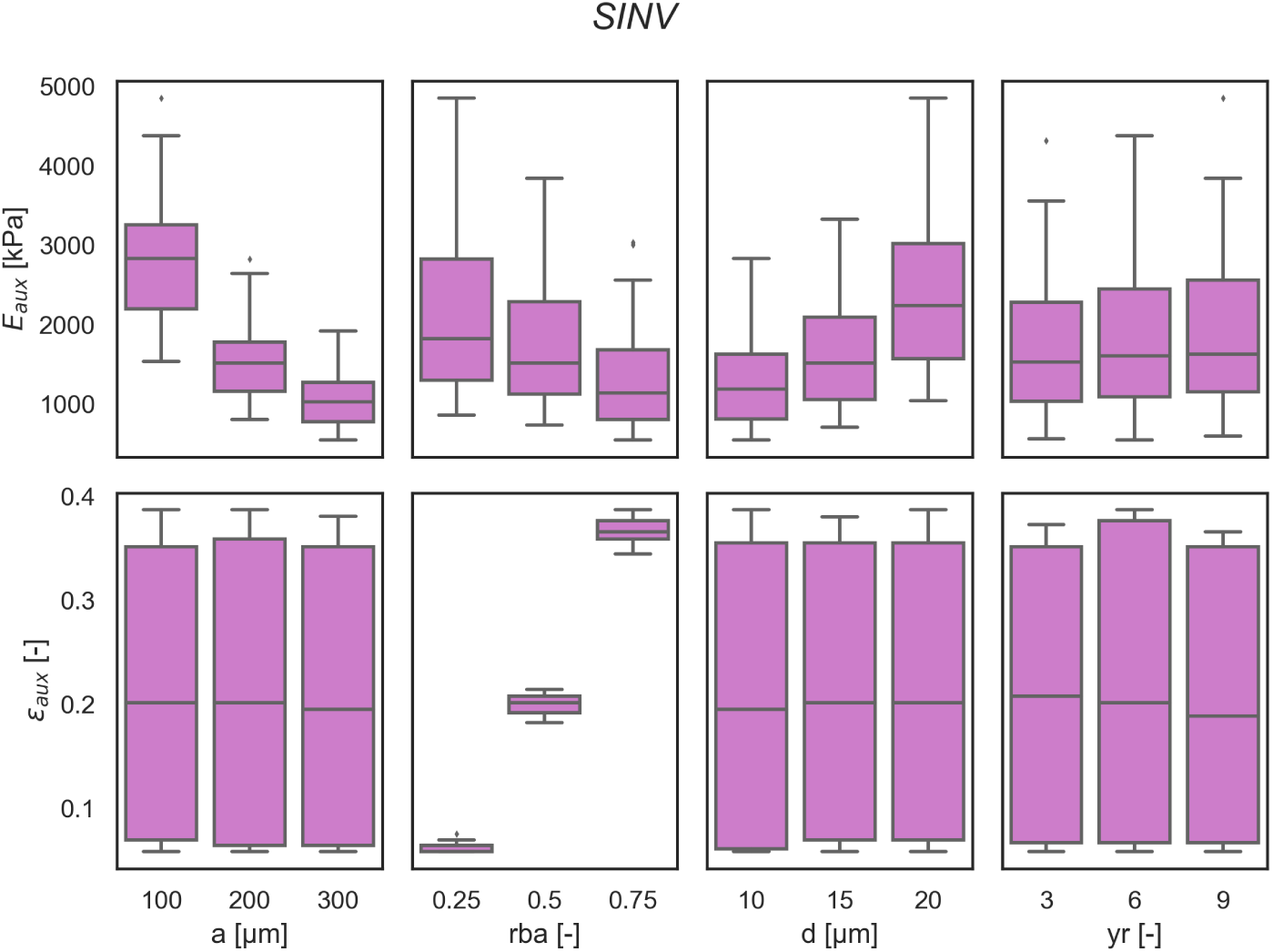
Interactions between geometric parameters and scaffold mechanical response variables observed from *SINV* design FEM simulations. *SINV* maintains low strain across all configurations while offering stiffness tunability, making it suitable for applications requiring limited deformability with mechanical control.

**Figure 11:**
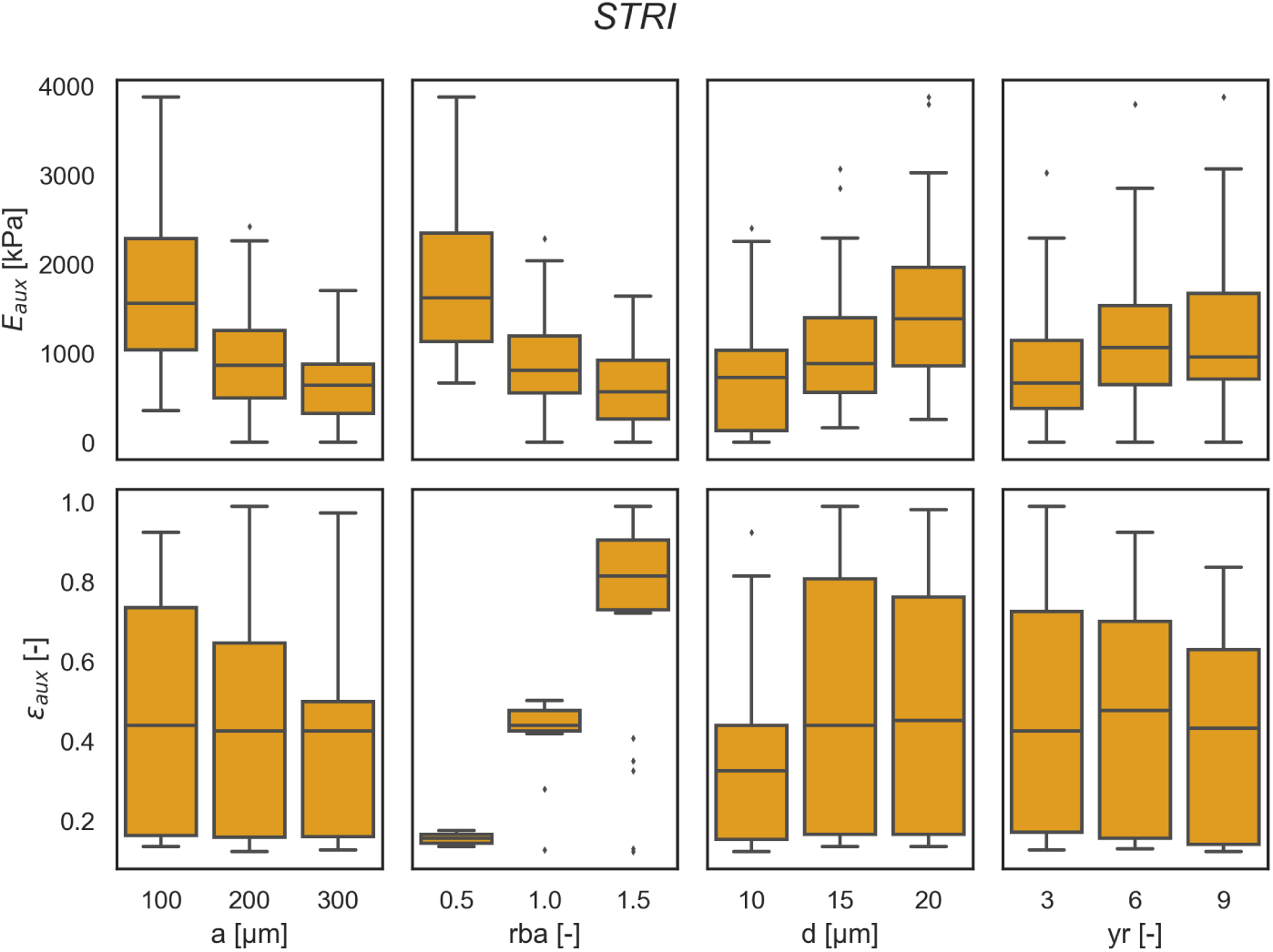
Interactions between geometric parameters and scaffold mechanical response variables observed from *STRI* design FEM simulations. *STRI* shows the greatest mechanical variability, with high sensitivity to multiple parameters, enabling flexible responses but reducing predictability.

### 8.2 Descriptive analysis of the FEM dataset

### 8.3 Sensitivity analysis of the *STRI* regression model

**Figure 12:**
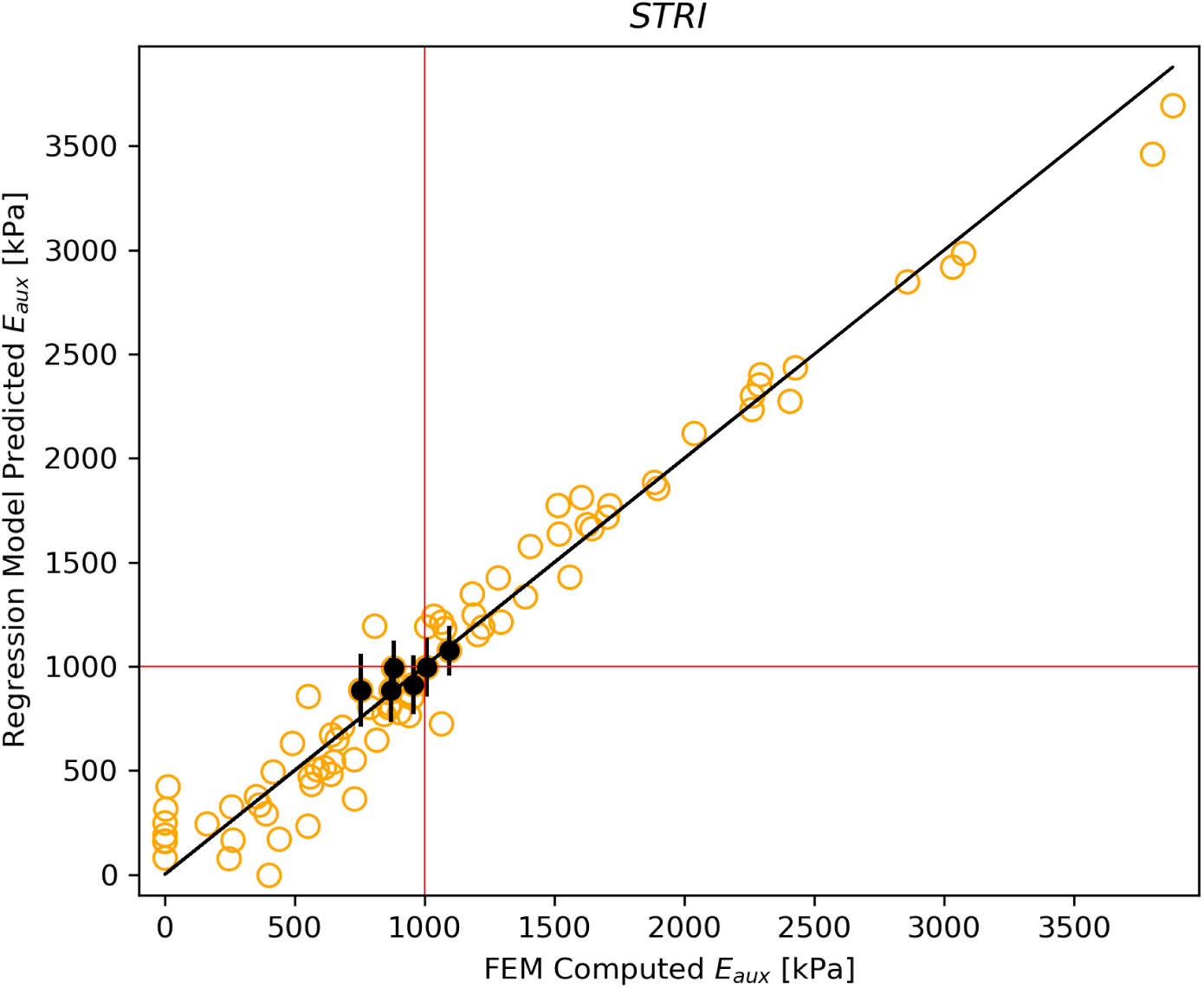
Predicted vs. FEM observed values of *E_aux_* for the *SINV* design, highlighting the predictions containing the target of 1000 kPa within their confidence interval of 95%.

### 8.4 Computational Tool Windows

**Table 2:**
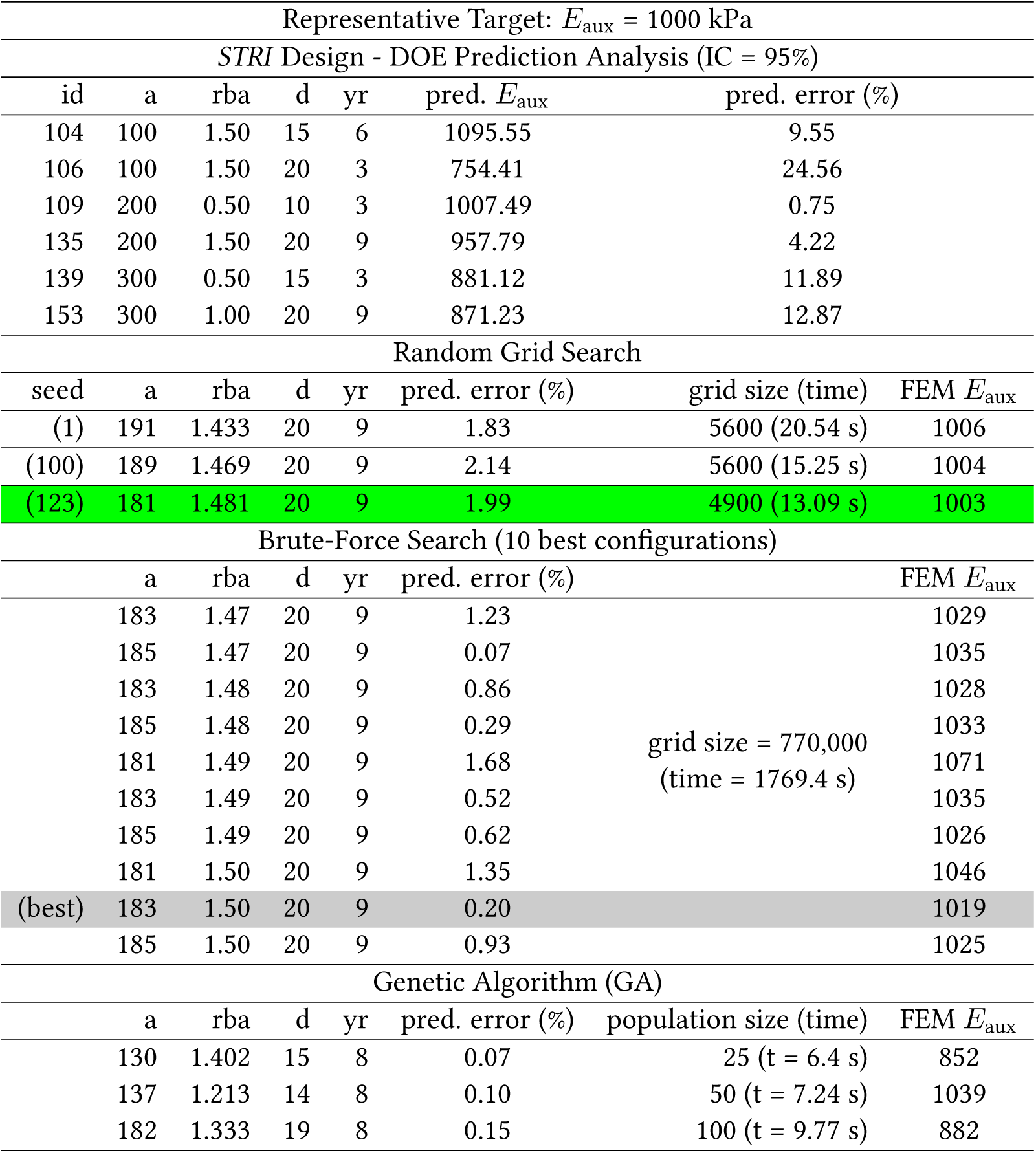
Summary of prediction performance across the different optimization methods tested. Although brute-force search identified the most accurate configuration (gray), random grid search achieved a similarly close solution (green) in significantly less time, regardless of the random seed used. This established random grid search as the most effective method. The genetic algorithm produced a comparable solution within a similar runtime but did not reach the same level of accuracy as the random grid approach.

**Figure 13:**
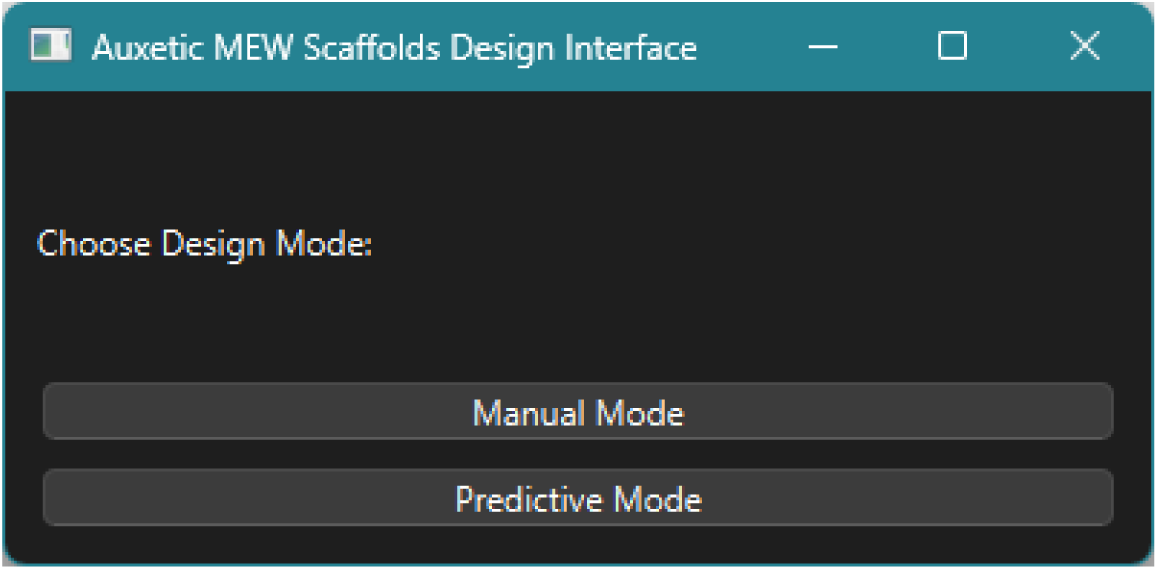
Main window of the computational tool, allowing to access the “Manual Mode” for direct design of prototypes, or “Predictive Mode” for target specification and best geometry selection.

**Figure 14:**
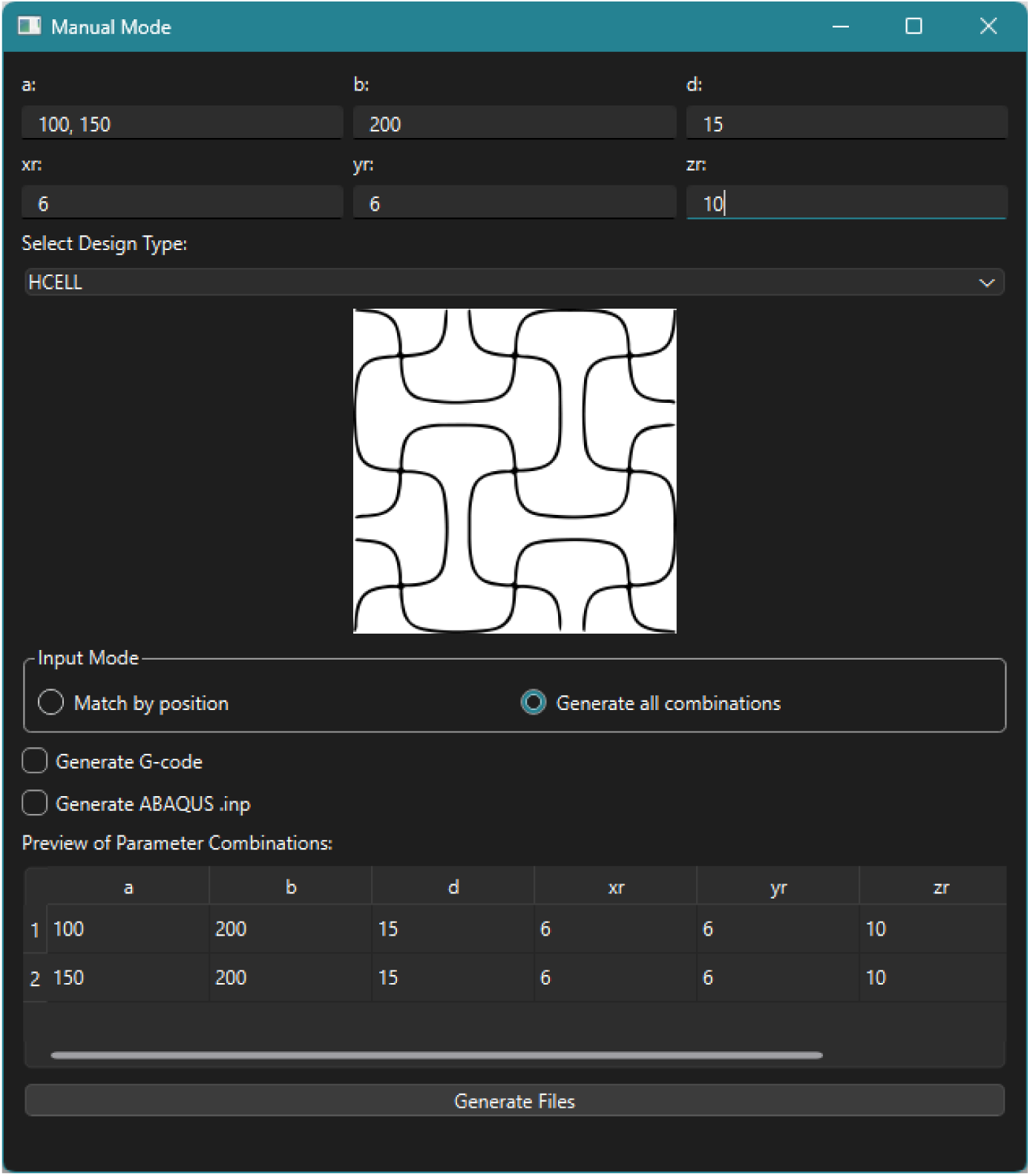
“Manual Mode” window of the computational tool. Includes some examples introduced to observe how introducing multiple values allows to generate the combinations of all of them. Additionally, individual configurations can be generated by introducing the same number of values for each parameter, where the position among the number of values will correspond to each individual configuration. By selecting any of the FEM or G-code boxes, it allows to export any or both files.

**Figure 15:**
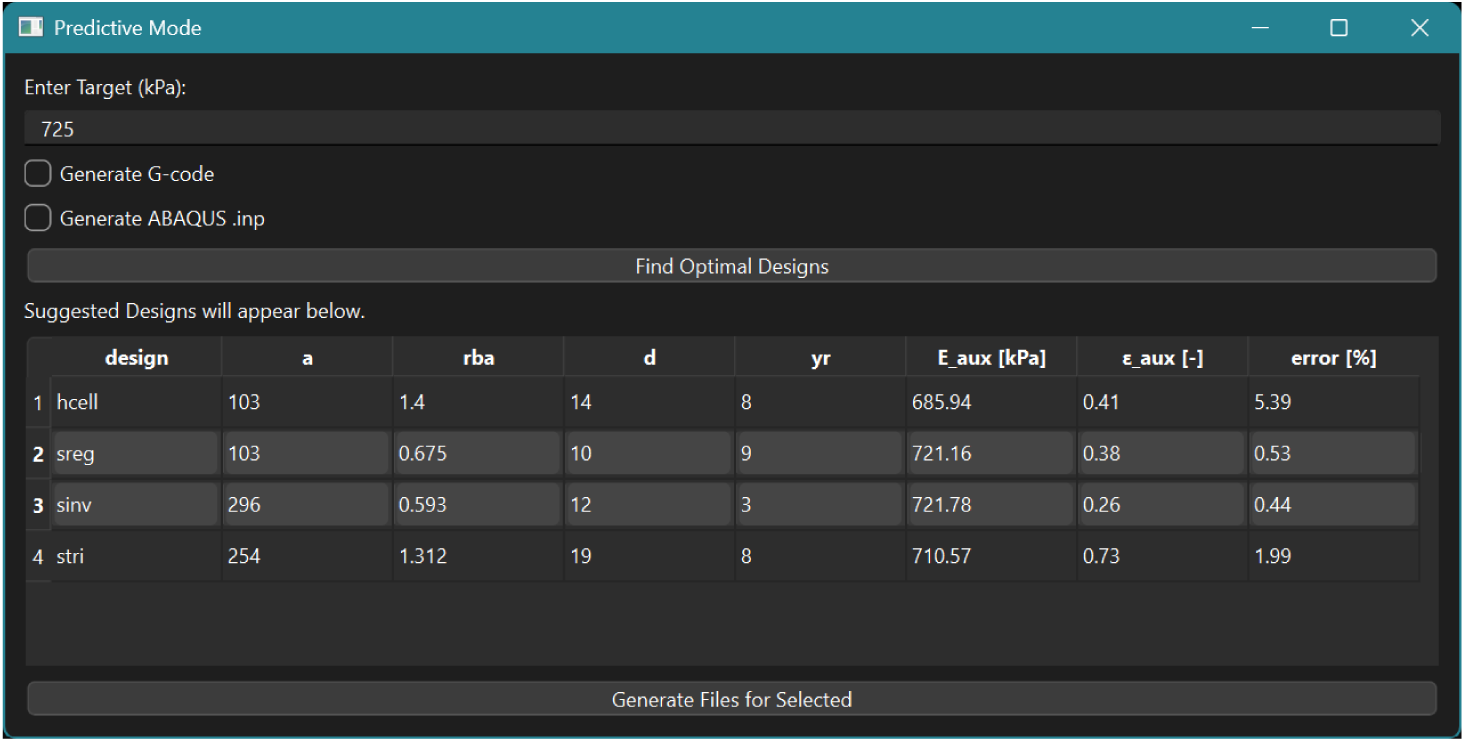
“Predictive Mode” window of the computational tool. Includes an example (the target of 725 kPa) to observe how the tool returns the best configuration found for each design and allows to evaluate them, select the desired options, and export the FEM files, G-code files, or both.

