## Supplementary material for "Regression-guided computational design of auxetic scaffolds for soft tissue applications": Supp. Material

### 1 Supplementary Material

#### 1.1 FEM simulation data post-processing

The post-processing of the FEM simulation data focused on extracting nodal displacements ($\Delta u$ [$\mu m$]) from the stretched edges and reaction forces ($RF$ [mN]) from the constrained edges, which allowed the calculation of effective stress ($\sigma$ [kPa]) and strain ($\varepsilon$ [$\mu m/\mu m$]) of the scaffolds following Eq. 1 and Eq. 2.

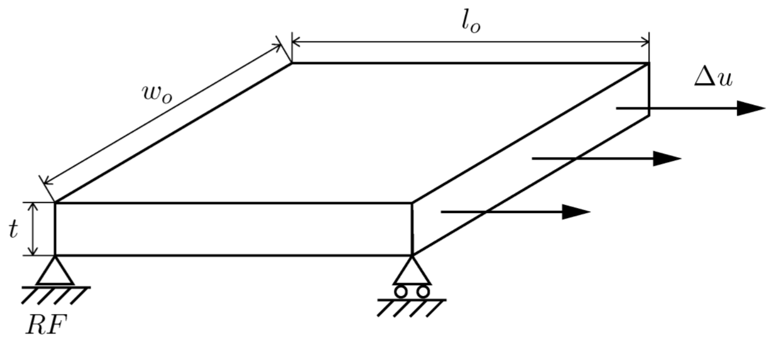


Figure S1: Schematic representation of the dimensions considered for scaffold mechanical properties calculation.

The scaffolds were treated as a homogeneous continuum, with their entire volume considered for length ($l_o$ [$\mu m$]), width ($w_o$ [$\mu m$]), thickness ($t$ [$\mu m$]), and the consequent transversal area ($A_t$ [$\mu m^2$]) determinations (Eq. 3), as illustrated in Figure S1.

$$\sigma = 10^6 * \frac{RF}{A_t} \tag{1}$$

$$\varepsilon = \frac{\Delta u}{l_o} \tag{2}$$

$$A_t = w_o * t \tag{3}$$

#### 1.2 Descriptive analysis of the FEM dataset

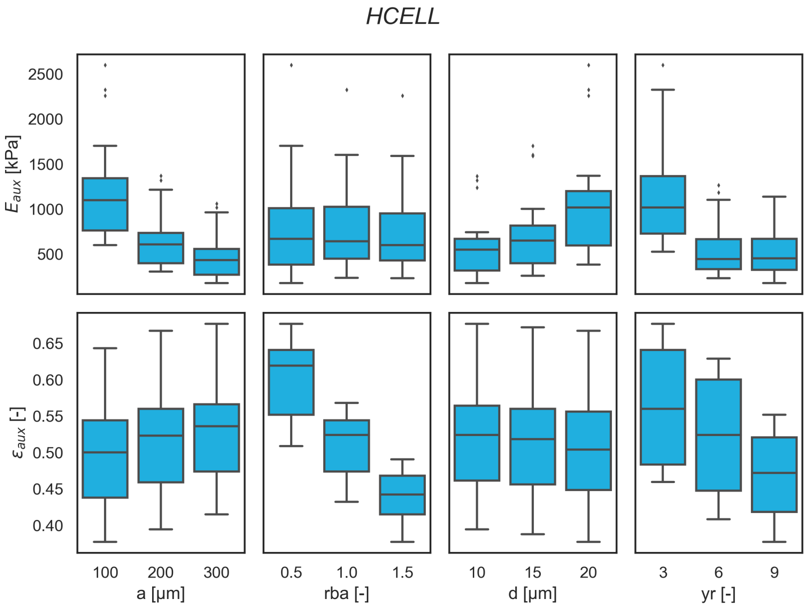


Figure S2: Interactions between geometric parameters and scaffold mechanical response variables observed from *HCELL* design FEM simulations. This design exhibits balanced sensitivity to all parameters, resulting in a stable and predictable mechanical response with moderate tunability in both stiffness and strain.

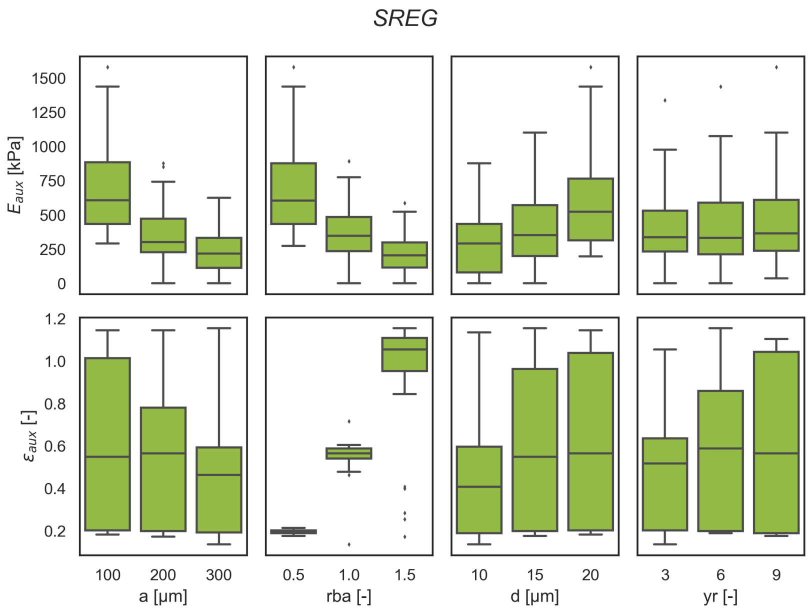


Figure S3: Interactions between geometric parameters and scaffold mechanical response variables observed from *SREG* design FEM simulations. Characterized by high strain variability and strong dependence on the "$rba$" ratio, *SREG* enables significant extensibility at the cost of increased mechanical unpredictability.

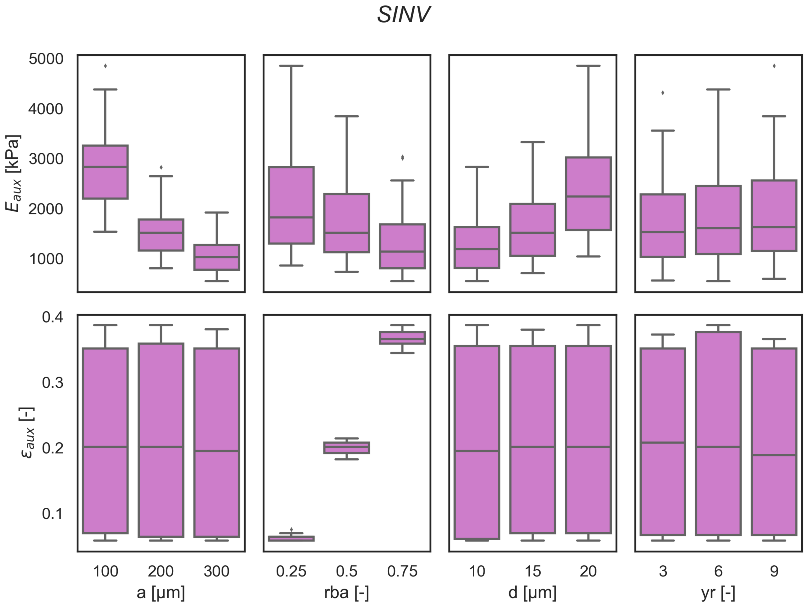


Figure S4: Interactions between geometric parameters and scaffold mechanical response variables observed from *SINV* design FEM simulations. *SINV* maintains low strain across all configurations while offering stiffness tunability, making it suitable for applications requiring limited deformability with mechanical control.

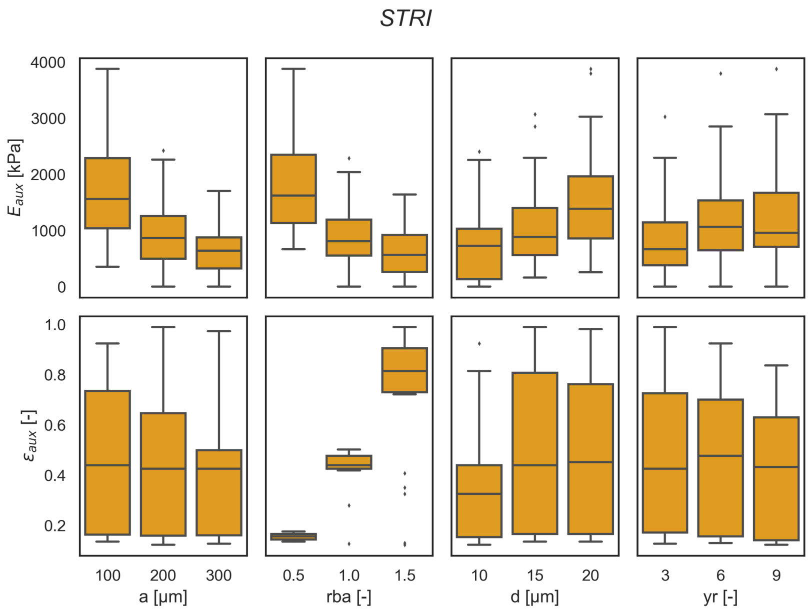


Figure S5: Interactions between geometric parameters and scaffold mechanical response variables observed from *STRI* design FEM simulations. *STRI* shows the greatest mechanical variability, with high sensitivity to multiple parameters, enabling flexible responses but reducing predictability.

#### 1.3 Sensitivity analysis of the *STRI* regression model

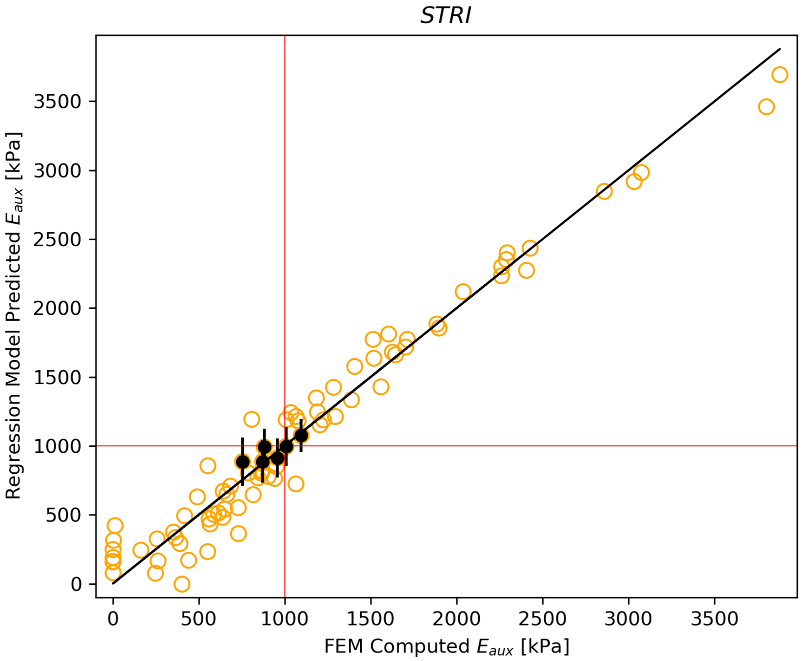


Figure S6: Predicted vs. FEM observed values of $E_{aux}$ for the *SINV* design, highlighting the predictions containing the target of 1000 kPa within their confidence interval of 95%.

| Representative Target: $E_{aux}$ = 1000 kPa | | | | | | | |
|---|---|---|---|---|---|---|---|
| *STRI* Design - DOE Prediction Analysis (IC = 95%) | | | | | | | |
| id | a | rba | d | yr | pred. $E_{aux}$ | pred. error (%) | |
| 104 | 100 | 1.50 | 15 | 6 | 1095.55 | 9.55 | |
| 106 | 100 | 1.50 | 20 | 3 | 754.41 | 24.56 | |
| 109 | 200 | 0.50 | 10 | 3 | 1007.49 | 0.75 | |
| 135 | 200 | 1.50 | 20 | 9 | 957.79 | 4.22 | |
| 139 | 300 | 0.50 | 15 | 3 | 881.12 | 11.89 | |
| 153 | 300 | 1.00 | 20 | 9 | 871.23 | 12.87 | |
| Random Grid Search | | | | | | | |
| seed | a | rba | d | yr | pred. error (%) | grid size (time) | FEM $E_{aux}$ |
| (1) | 191 | 1.433 | 20 | 9 | 1.83 | 5600 (20.54 s) | 1006 |
| (100) | 189 | 1.469 | 20 | 9 | 2.14 | 5600 (15.25 s) | 1004 |
| (123) | 181 | 1.481 | 20 | 9 | 1.99 | 4900 (13.09 s) | 1003 |
| Brute-Force Search (10 best configurations) | | | | | | | |
| | a | rba | d | yr | pred. error (%) | | FEM $E_{aux}$ |
| | 183 | 1.47 | 20 | 9 | 1.23 | grid size = 770,000 (time = 1769.4 s) | 1029 |
| | 185 | 1.47 | 20 | 9 | 0.07 | | 1035 |
| | 183 | 1.48 | 20 | 9 | 0.86 | | 1028 |
| | 185 | 1.48 | 20 | 9 | 0.29 | | 1033 |
| | 181 | 1.49 | 20 | 9 | 1.68 | | 1071 |
| | 183 | 1.49 | 20 | 9 | 0.52 | | 1035 |
| | 185 | 1.49 | 20 | 9 | 0.62 | | 1026 |
| | 181 | 1.50 | 20 | 9 | 1.35 | | 1046 |
| (best) | 183 | 1.50 | 20 | 9 | 0.20 | | 1019 |
| | 185 | 1.50 | 20 | 9 | 0.93 | | 1025 |
| Genetic Algorithm (GA) | | | | | | | |
| | a | rba | d | yr | pred. error (%) | population size (time) | FEM $E_{aux}$ |
| | 130 | 1.402 | 15 | 8 | 0.07 | 25 (t = 6.4 s) | 852 |
| | 137 | 1.213 | 14 | 8 | 0.10 | 50 (t = 7.24 s) | 1039 |
| | 182 | 1.333 | 19 | 8 | 0.15 | 100 (t = 9.77 s) | 882 |

Table S1: Summary of prediction performance across the different optimization methods tested. Although brute-force search identified the most accurate configuration (gray), random grid search achieved a similarly close solution (green) in significantly less time, regardless of the random seed used. This established random grid search as the most effective method. The genetic algorithm produced a comparable solution within a similar runtime but did not reach the same level of accuracy as the random grid approach.

#### 1.4 Computational Tool Interface

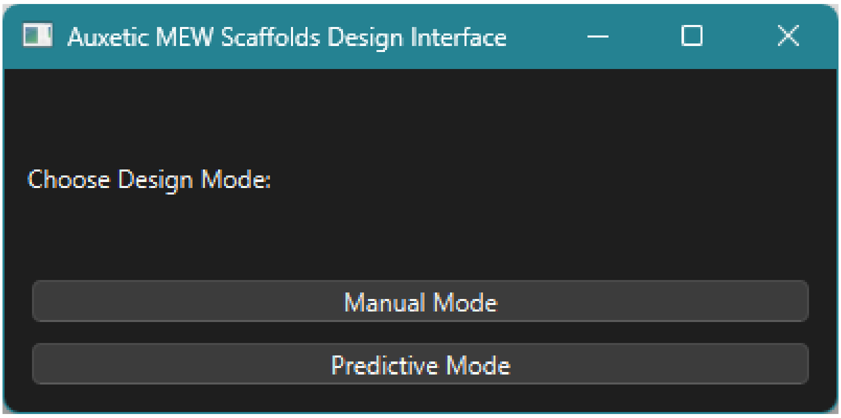


Figure S7: Main window of the computational tool, allowing to access the "Manual Mode" for direct design of prototypes, or "Predictive Mode" for target specification and best geometry selection.

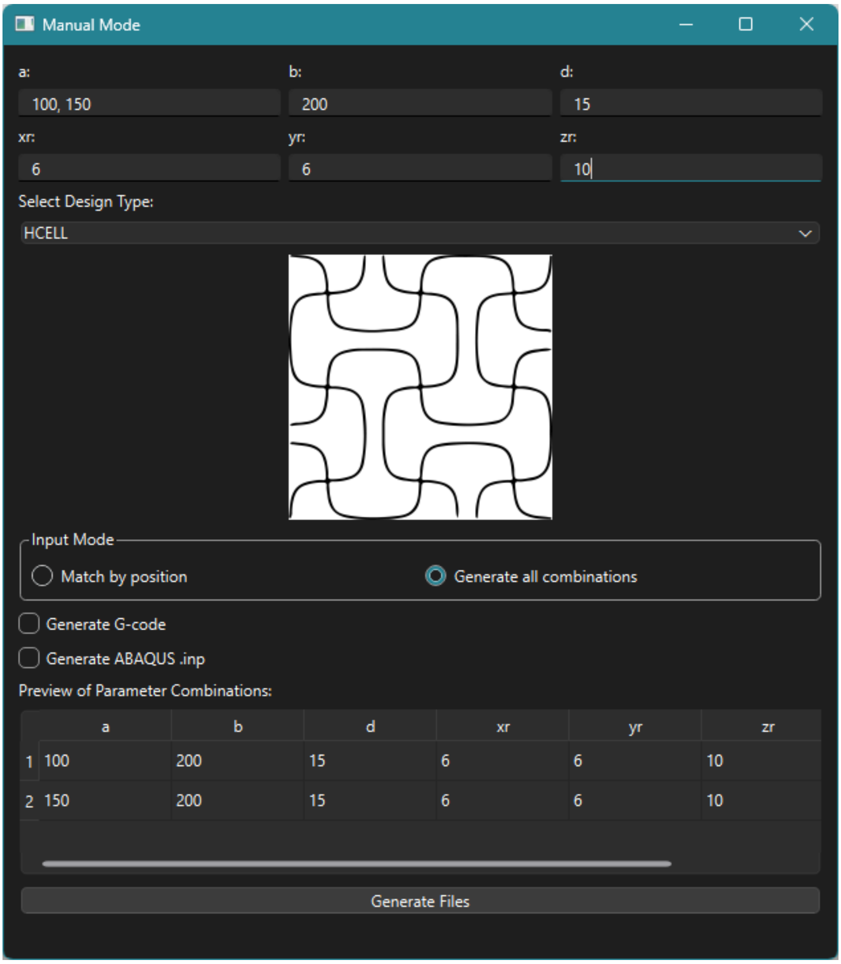


Figure S8: "Manual Mode" window of the computational tool. Includes some examples introduced to observe how introducing multiple values allows to generate the combinations of all of them. Additionally, individual configurations can be generated by introducing the same number of values for each parameter, where the position among the number of values will correspond to each individual configuration. By selecting any of the FEM or G-code boxes, it allows to export any or both files.

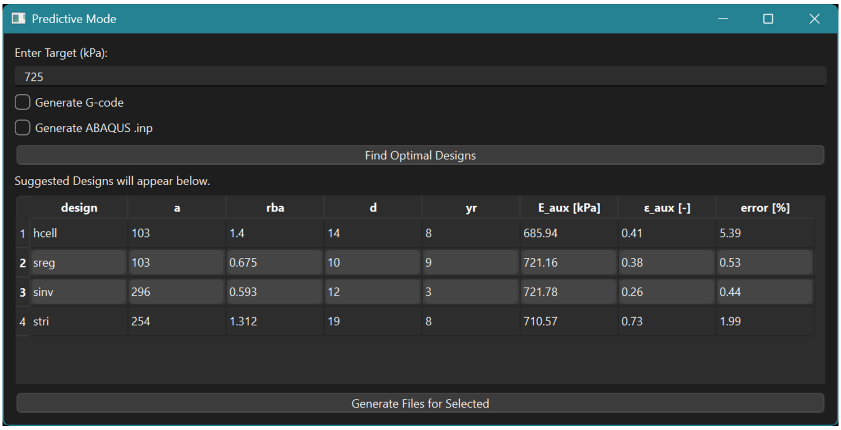


Figure S9: "Predictive Mode" window of the computational tool. Includes an example (the target of 725 kPa) to observe how the tool returns the best configuration found for each design and allows to evaluate them, select the desired options, and export the FEM files, G-code files, or both.
